# A hippocampal digital Twin reveals latent structure in freely flying bats

**DOI:** 10.64898/2026.10.09.758148

**Authors:** Adam S. Lowet, Yongrong Qiu, Angelo Forli, Krishna Chaitanya Kasuba, Yang Lin, Andreas S. Tolias, Michael M. Yartsev

## Abstract

Concurrent advances in artificial intelligence (AI), large-scale neuronal recording, and behavioral tracking have enabled increasingly powerful predictive models of brain-behavior relationships. Yet predictive accuracy does not automatically yield interpretable principles of neural organization, particularly during natural behavior, where the relevant variables computed are often unknown in advance. Here, we developed a functional digital twin of the hippocampus, trained jointly on large-scale neural activity and self-directed three-dimensional flight in freely flying bats. A multi-modal, multitask transformer accurately predicted both neural activity and behavior, outperformed standard approaches, and generalized across recording sessions. Interrogating the model’s representations suggested a form of hippocampal generalization across self-selected trajectories that was not apparent in physical coordinates, but was plainly visible in the model’s low-dimensional latent space. We confirmed this organization directly in the recorded neural activity: neurons fired at corresponding stages of spatially distinct paths connecting the same origin and destination, preserving their relative firing order even as their firing fields shifted with the path. Analysis of neuron-specific readouts further identified functional relationships among neurons that generalized from flight to waking rest and sleep, brain states never used for training, and differed systematically between the hippocampal subregions CA1 and CA3. Together, these results show how AI-based functional digital twins can complement conventional analysis by generating testable hypotheses about neural organization during natural behavior.

## 2 Introduction

A central goal in neuroscience is to understand how neural activity gives rise to cognition as animals interact naturally with their environment. Addressing this goal often requires naturalistic paradigms that preserve the complexity and self-directed structure of behavior [1]. Recent advances in large-scale neuronal recording [2–4] and high-resolution behavioral tracking [5–7] can now capture this complexity at unprecedented scale. Yet extracting organizing principles from these data remains challenging. Controlled experiments often isolate selected sensory or behavioral variables to test specific hypotheses, whereas natural behavior involves many interacting variables whose relevance to neural activity may be unknown in advance.

Numerous computational approaches have emerged to address this challenge [8]. Models that prioritize simplicity, interpretability, or mathematical tractability provide useful accounts of neural activity, but their assumptions about neural dynamics or relevant behavioral variables can constrain the structure they reveal. Large-scale machine learning methods offer a more flexible approach to modeling these complex relationships, at the expense of analytic tractability. These methods have transformed fields ranging from vision and language to protein folding [9–11] and are increasingly being applied to jointly model neural and behavioral data [12–21]. Recent neural foundation models trained across animals and recording sessions can generalize to new subjects and stimulus domains and can recover biological properties that were not explicitly provided during training [21].

These advances enable the construction of functional “digital Twins”: predictive models of biological systems that support large-scale virtual experiments, gradient-based analyses of predicted neural responses, and interrogation of internal representations and parameters to reveal principles of neural organization. However, functional digital twins have primarily been applied in sensory systems under constrained, often head-fixed paradigms, in which the experimenter — not the ani-mal — defines the inputs [13,20–27]. How well these models capture activity in circuits supporting memory and navigation, and whether interrogating them can expose biologically meaningful latent structure, remain open questions — particularly in freely moving animals engaged in spontaneous and self-directed natural behavior.

The mammalian hippocampus, which plays an essential role in spatial memory and navigation [28], provides a compelling test bed for this approach. Spontaneous navigation often lacks an experimenter-defined trial structure, and the organizing principles of hippocampal activity remain abstract and debated [29–31]. Hippocampal place cells — best known for their spatial selectivity [32,33] — are highly sensitive to behavioral context, often distinguishing similar locations according to direction of travel, recent experience, anticipated future behavior, or internal goals [34–48]. At the same time, theoretical accounts emphasize the hippocampus’ capacity to generalize relational structure across experiences [49,50], including in non-spatial domains [51–60]. Experimental studies have tended to identify these distinctions and generalizations by testing hypotheses about specific behavioral variables chosen in advance. Whether such organizing structure can instead be inferred directly from large-scale neural and behavioral data — and thereby reveal unexpected structure within the hippocampal cognitive map — remains unresolved.

Here we developed a multimodal, multitask transformer [19,20,61] that jointly models hippocampal activity and three-dimensional navigation in freely flying Egyptian fruit bats (*Rousettus aegyptiacus*) navigating in the lab and in an outdoor field station [45]. Trained on data from 3,080 neurons across 1,738 flights, 17 recording sessions and six animals, the model accurately predicted single-neuron activity, decoded flight trajectories, and generalized to held-out sessions. We then interrogated its latent representations and neuron-specific readouts to generate testable biological hypotheses about the underlying hippocampal system. The digital Twin’s latent space organized diverse 3D flight paths into recurring directional motifs that generalized across animals and routes. This organization predicted that neurons should fire at corresponding stages of spatially distinct paths, preserving relative sequence order despite shifts in their physical firing fields; we confirmed this prediction directly in the recorded neural activity. Neuron-specific readouts further captured functional relationships that generalized from flight to waking rest and sleep — brain states never used to train the model — and differed systematically between CA1 and CA3. Together, these results show how functional digital Twins can move beyond prediction to generate and test hypotheses about neural organization during natural behavior.

## 3 Results

### 3.1 A functional digital Twin achieves superior prediction of hippocampal activity and behavior

We trained a deep neural network that jointly models binned spike counts and time-aligned three-dimensional position. During training, half of the neural and behavioral tokens were masked on each forward pass, and the model was trained to reconstruct the missing values (Fig. S1A–B; see Methods). This multimodal, multitask architecture enabled a unified model to perform behavioral decoding (predicting behavior from neural activity), neural encoding (predicting neural activity from behavior), and any combination thereof [19,20], all with a single set of parameters (Fig. S1C, Table S1). A shared Transformer core mapped data from different animals and sessions into a common latent space, while session-and environment-specific linear mappings projected neural activity and behavior into and out of this space (Fig. 1A). We subsequently interrogated the latent embeddings to characterize population-level neural organization (Figs. 2–3) and the neuron-specific readout weights to relate model structure to anatomical and functional properties measured *in vivo* (Fig. 4).

**Fig. 1.**
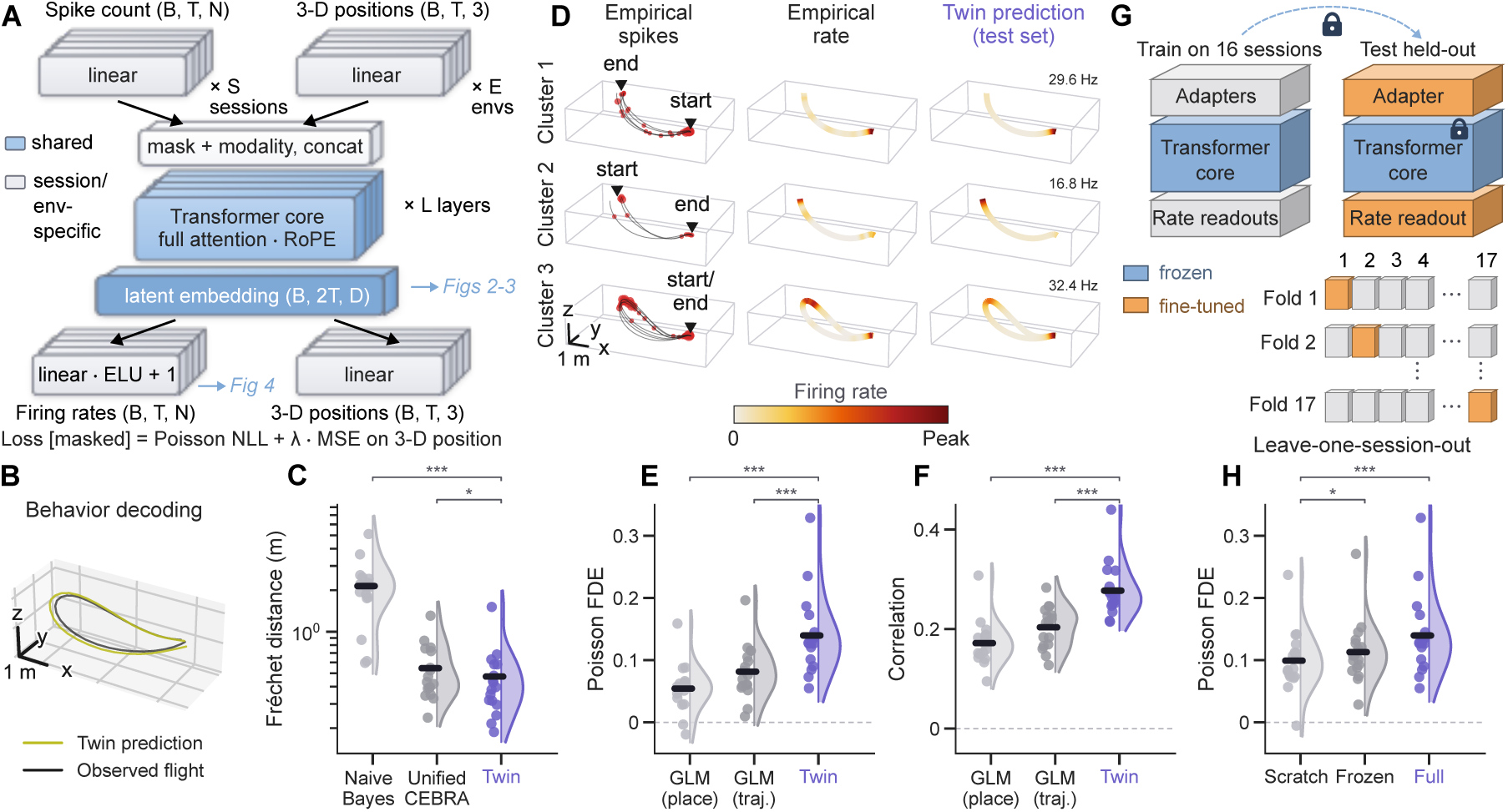
A functional digital Twin achieves superior prediction of hippocampal activity and behavior. (A) Twin model architecture. Subsequent figures analyze particular components of the model, specifically the latent embeddings (Figs. 2–3) and neural readout weights (Fig. 4). *B* = batch size = number of flights per session, *T* = sequence length = number of time bins, *N* = number of neurons per session, *S* = number of sessions, *E* = number of environments, *L* = number of transformer layers, *D* = transformer latent dimension, *ELU* = Exponential Linear Unit nonlinearity, *NLL* = negative log likelihood, *λ* = decoding weight, *MSE* = mean squared error. For further details see Methods. (B) Representative decoded (olive) versus observed (black) 3-D flight trajectory from the Twin decoder. (C) Decoding accuracy as per-flight 3-D discrete Fŕechet distance (lower is better) for a ‘Näıve Bayes’ place decoder, ‘Unified CEBRA’ decoder, and the digital Twin model. Here and for all similar plots throughout the paper, dots show individual sessions (per-session medians), the half-violin their distribution, and the thick ink bar the across-session mean, with the model of interest in purple; *n* = 17 sessions. Wilcoxon vs. the Twin: ‘Näıve Bayes’ (*P* = 1.53 *×* 10*^−^*^5^), ‘Unified CEBRA’ (*P* = 0.0448). (D) An example unit with spatially separated place fields on three different flight clusters (*rows*). Flight start and end are indicated. *Left*, observed spikes (red dots, area *∝* binned spike count) on the black flight paths; *center*, mean empirical firing rate map; *right*, the Twin’s mean predicted rate map. Within each row, the empirical and predicted maps share one color scale, normalized to the larger of their two peaks; *inset*, peak rate is indicated for each empirical and predicted pair. (E–F) Encoding fit quantified as per-neuron Poisson fraction deviance explained (FDE, E) and Pearson correlation (F) between predicted and observed spike trains; Wilcoxon vs. the Twin: (E) ‘place GLM’ (*P* = 1.53 *×* 10*^−^*^5^), ‘trajectory-specific GLM’ (*P* = 3.05 *×* 10*^−^*^5^);(F) ‘place GLM’ and ‘trajectory-specific GLM’ both *P* = 1.53 *×* 10*^−^*^5^. (G) Schematic of transfer learning analysis. Model architecture is identical to (A). All model weights were frozen (padlocks, blue) except for the session-specific neural readout weights, which were fine-tuned (orange). For each held-out session, a new base model was trained and then frozen as in (G) before fine-tuning the readouts on the held-out session (orange card). (H) Poisson FDE for the Twin, either retrained from scratch on a single session, transferred to the held-out session with readout fine-tuning only (G), or evaluated on the full 17-session model (E). Wilcoxon vs. from-scratch: ‘frozen’ *P* = 0.0174 and ‘full’ *P* = 2.90 *×* 10*^−^*^4^. Here and for all figures, *^∗^P <* 0.05, *^∗∗^P <* 0.01, *^∗∗∗^P <* 0.001; n.s. comparisons are generally unmarked. *N* = 17 sessions from 6 bats (4 bats in an indoor flight room and 2 bats in an outdoor field station, 54-394 neurons per session, 62-145 flights per session); decoding and encoding performance is measured on a held-out test set not used for fitting or hyperparameter selection. Unless otherwise indicated, data is colored by bat consistently across figures. See also Figs. S1–S3.

**Fig. 2.**
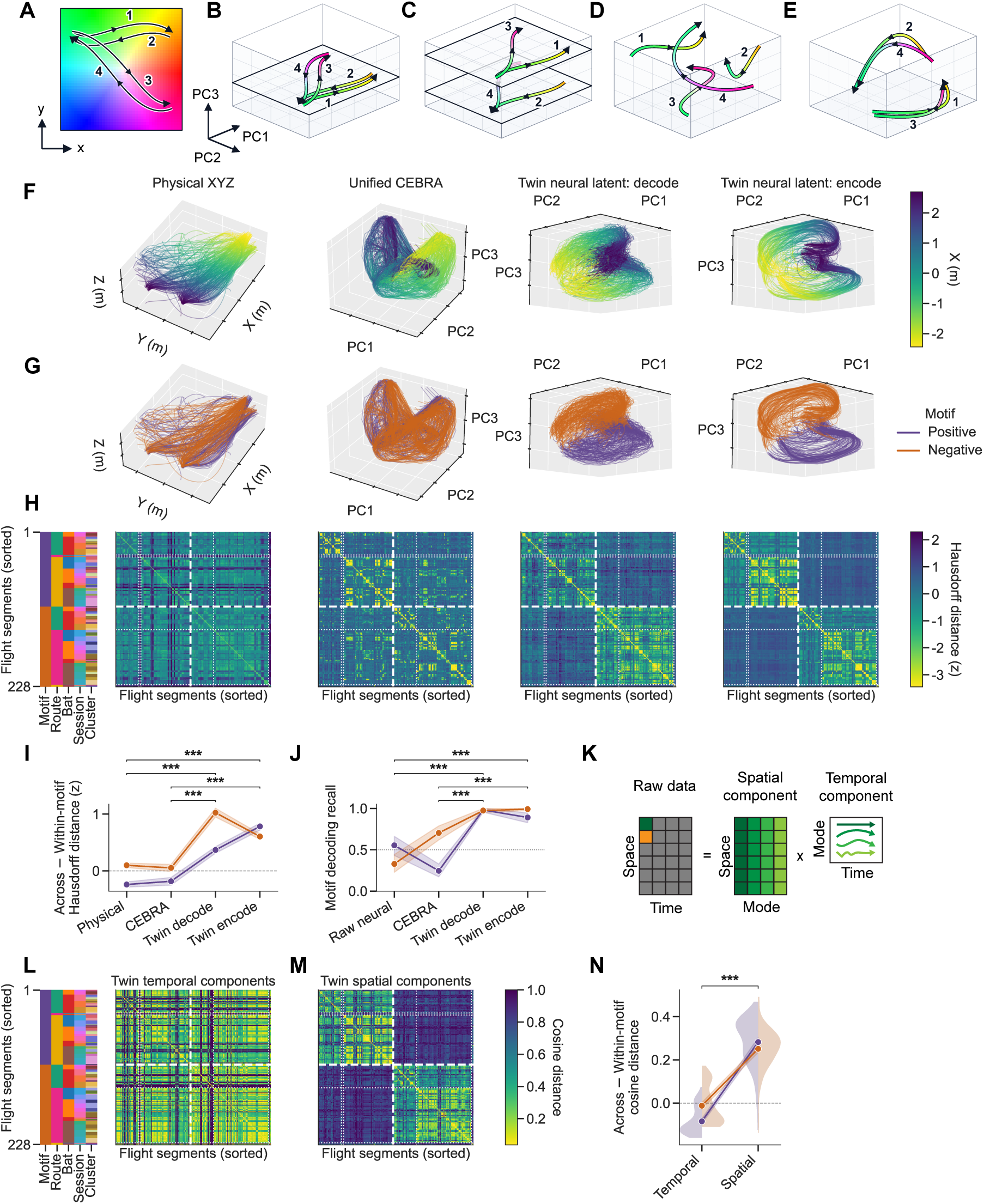
The Twin latent representation warps physical space to align geometry and dynamics across animals. (A–E) Schematics of four flight trajectories (labeled 1–4) in physical space (A) and those same four flights in four different, hypothetical latent space representations (B–E). (A) Top-down (x-y) projection of flights (directed arrows) in physical space. The color map represents a smooth Euclidean metric over physical space, and is consistent in panels B–E. (B) One hypothetical neural code for the four flights in (A), in which the neural space is a faithful Euclidean representation of physical space. Like colors (physical locations) therefore map to identical points in neural space. (C) Same as (B), except that flights with different x-directions (arrows) are projected onto distinct neural manifolds along the PC3 dimension. This is the code implied by direction-specific place cells with a single dominant direction of travel [34–36]. (D) Individual flight trajectories correspond to continuous progressions through the neural space, but the relationship between distinct trajectory types is random, as implied by cluster-specific place cells [42]. Similar physical locations (colors) on different flights are therefore scattered throughout the neural space. (E) Different x-directions cluster into distinct lobes in the neural space, while other spatial dimensions are collapsed, indicating a profound degree of spatial generalization. Compare to the rightmost panels of F–G. (F–H) Four representations are compared column-by-column throughout: true 3-D position (Physical XYZ), the ‘Unified CEBRA’ latent, and the digital Twin 64-D neural latent generated in decode mode and encode mode. (F) Held-out flights, colored by the bat’s instantaneous x-coordinate (*n* = 288 segments from all 232 test flights). To obtain the three-dimensional representations, raw 64-dimensional CEBRA and Twin embeddings were projected using PCA. For Twin, a single shared PCA basis was fit jointly across the encode and decode latents, whereas a distinct PCA basis was fit independently for CEBRA. (G) Same as (F), but colored by motif (flying in the positive or negative principal direction). Unlike physical space and the CEBRA embedding, the Twin latent segregates segments by motif and not simply by position. (H) Representational similarity analysis (RSA): every pair of *n* = 228 segments (190 test flights) that included reward as start or end is compared by the symmetric Hausdorff distance between their point clouds in physical or raw latent space and converted to a per-space z-score, then hierarchically sorted by motif, route, bat, session and cluster (barber pole plot, *left* ; motif colored as in (G), route colored such that reward*→*reward is green, reward*→*perch magenta and perch*→*reward gold). Thick white dashed lines mark the motif boundary, thin dotted lines the route boundaries within each motif. (I) Across *−* Within-motif Hausdorff distance per segment (mean *±* 95% CI band, colored by motif as in G; dashed line at zero), counting only partners of a *different* route. Two-sided Wilcoxon signed-rank tests compare the Twin decode and encode latents against Physical XYZ (*P* = 1.62 *×* 10*^−^*^37^ decode, *P* = 4.14 *×* 10*^−^*^38^ encode) and ‘Unified CEBRA’ (*P* = 5.78 *×* 10*^−^*^36^ decode, *P* = 5.88 *×* 10*^−^*^38^ encode). (J) Motif decoded from the time-averaged segment representation using a support vector classifier with radial basis function kernel, by leave-one-route-out: each route in turn is held out and predicted by a classifier trained only on the other routes, so every segment is held out exactly once and nothing about route identity carries from training to test. The two lines are the per-motif recalls, whose mean is the balanced accuracy (chance = 0.5, dotted line). “Raw neural” fits a new segment-wise decoder for each session, based on time-averaged spike counts. McNemar tests compare the Twin decode and encode latents against ‘Raw neural’ (*P* = 2.14 *×* 10*^−^*^17^ decode, *P* = 4.10 *×* 10*^−^*^15^ encode) and ‘Unified CEBRA’ (*P* = 4.00 *×* 10*^−^*^25^ decode, *P* = 8.82 *×* 10*^−^*^24^ encode). (K) Schematic of dynamic mode decomposition (DMD): a segment’s latent time series is factored into a product of spatial modes and their temporal dynamics, each mode oscillating at a single frequency with its own growth or decay rate. (L) RSA on the eigenvalue (temporal) profile of each motif segment in the Twin encode latent space, sorted and arranged as in (H). (M) Same as (L), but RSA on the real part of the (spatial) DMD mode with the largest power, taken after purely real (non-oscillatory) modes are dropped and each complex-conjugate pair is consumed, so the ranking counts distinct oscillations. (N) Across *−* Within-motif cosine distance from (L) and (M), again over cross-route partners only; split violins show the per-segment distribution behind each mean *±* 95% CI. Motif is carried far more strongly by the spatial modes than by the eigenvalue spectra (*P* = 2.95 *×* 10*^−^*^38^, *n* = 228 segments). See also Figs. S4–S7.

**Fig. 3.**
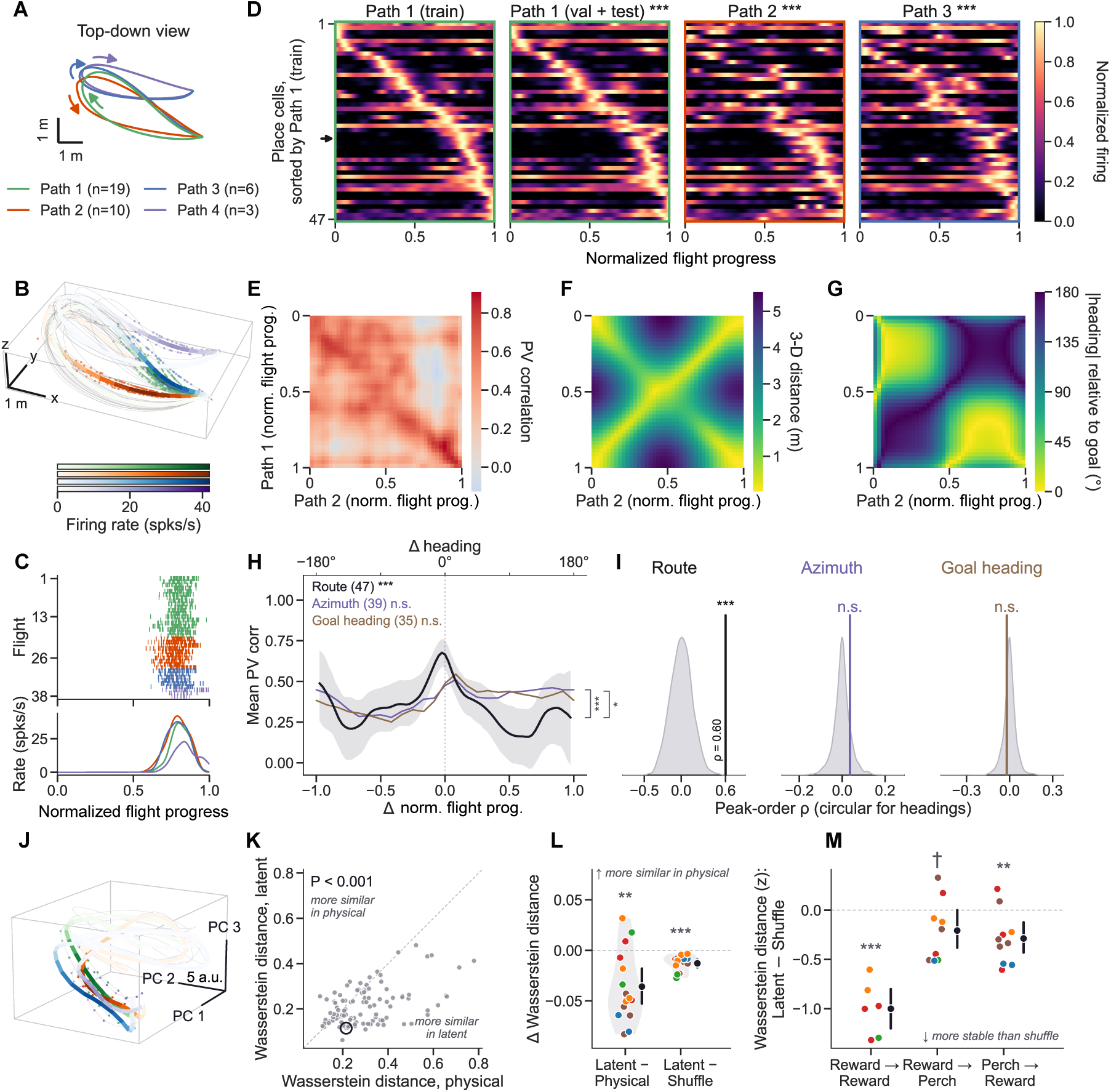
Latent representations expose spatial generalization at the single-neuron and population level. (A) Top-down (x–y) view of the four example mean flight paths — Path 1 (green), Path 2 (orange), Path 3 (blue) and Path 4 (purple), all belonging to one reward *→* reward route of a single session — with arrows indicating travel direction. (B) Rate map of an example unit: its firing rate (color) drawn along the four mean flight paths in 3-D (Greens/Oranges/Blues/Purples), with the individual flights faint underneath and individual spikes as dots. The unit fires differently on the four paths as a function of spatial position. (C) Per-flight normalized-flight-progress spike raster (*top*) and mean firing-rate tuning curves (*bottom*) for the same unit on each path reveal that it fires at a similar normalized flight progress on all of them despite their spatial divergence. (D) Firing-rate sequences of all place cells (*n* = 47 place cells) in normalized flight progress on the Path 1 training flights, the held-out Path 1 validation + test flights, all Path 2 flights and all Path 3 flights (*left* to *right*, all paths with *n ≥* 5 flights). Each cell’s rate is normalized to its own peak and cells are sorted by their peak progress on the Path 1 training flights. Each panel is framed in its path’s color. The peak-progress order is highly preserved between the Path 1 training and held-out flights (Spearman *ρ* = 0.909, *P* = 5.00 *×* 10*^−^*^4^), and also between the Path 1 training flights and Path 2 (*ρ* = 0.534, *P* = 5.00 *×* 10*^−^*^4^) and Path 3 (*ρ* = 0.543, *P* = 5.00 *×* 10*^−^*^4^). The arrowhead marks the example unit in (B, C, J). (E) Population-vector (PV) correlation between Path 1 (rows) and Path 2 (columns) across normalized-flight-progress bins shows strong correlations along the main diagonal. (F) 3-D Euclidean distance between the Path 1 and Path 2 mean positions at each pair of normalized-flight-progress bins. Note the strong anti-diagonal band, which is not present in the PV correlation matrix, showing a dissociation between spatial position and population coding. (G) Absolute difference in goal heading — flight heading taken relative to the bearing to the reward feeder each flight lands on — between the Path 1 and Path 2 mean paths. Note the block diagonal structure which is similarly absent from (E). (H) Mean PV correlation along each diagonal versus lag (normalized flight progress, black, with 95% bands from bootstrapping over place cells), and the same curve after re-indexing the PV matrix on allocentric azimuth or on goal heading, each computed on its own cross-validated heading-tuned cells (39 azimuth-, 35 goal-heading-tuned; the top axis reads heading lag in degrees). Key entries carry each curve’s near-versus-far test. Near = within *±*10% of that axis’s range, far = *≥* 50% of it, against a circular-shift permutation null: normalized flight progress +0.321 (*P* = 5.00*×*10*^−^*^4^), azimuth +0.066 (*P* = 0.39), goal heading +0.102 (*P* = 0.090). Brackets are paired cell-bootstrap comparisons of the normalized-flight-progress curve against each heading curve (normalized flight progress *−* azimuth Δ = +0.203, *P* = 1.00 *×* 10*^−^*^3^; *−* goal heading Δ = +0.125, *P* = 0.019). (I) Distribution of the peak-order correlation between the held-out Path 1 flights and Path 2 under a peak-order shuffle null (gray), with the observed value marked, for each of the three axes of (H): normalized flight progress (Spearman *ρ* = 0.601, *P* = 5.00 *×* 10*^−^*^4^), azimuth (*ρ* = 0.036, *P* = 0.14) and goal heading (*ρ* = *−*0.019, *P* = 0.75). (J) The example unit’s rate map in the Twin latent space, on the rate scale of (B). Only test-split flights carry a latent (*n* = 2 / 3 / 1 / 2 flights for Paths 1–4). (K) Per-neuron Wasserstein distance between the rate maps of the different paths of a route (normalized to the longest transport the bin grid admits, so lower = more stable) in the latent versus the physical space, for every neuron and route of the example session (*n* = 108 neuron–route pairs; Wilcoxon signed-rank *P* = 4.4 *×* 10*^−^*^9^). The dashed line is unity and the circled point is the unit of (B, C, J) on its reward *→* reward route. (L) Wasserstein distance differences pooled over routes, one point per session colored by bat, with mean *±* 95% CI: latent *−* physical (Wasserstein *P* = 2.6 *×* 10*^−^*^3^) and latent *−* occupancy-preserving circular-shift shuffle (Wasserstein *P* = 6.1 *×* 10*^−^*^5^), session-level Wilcoxon over *n* = 15 sessions; negative values mean the latent rate map is the more stable. (M) Bin-free Wasserstein distance of the 15-D latent, expressed as a *z* score against its own circular-shift null (latent *−* shuffle, in units of the null’s SD; negative = more stable than shuffle), by route: reward *→* reward −0.99 (*n* = 6 sessions, *P* = 3.0 *×* 10*^−^*^4^), reward *→* perch −0.19 (*n* = 9, *P* = 0.083), perch *→* reward −0.32 (*n* = 10, *P* = 8.1 *×* 10*^−^*^3^); (M) uses one-sample *t*-tests on the session medians rather than the Wilcoxon signed-rank of (L), as too few sessions contribute to each route for the signed-rank to resolve these *P* values. This estimator puts no grid on either space, which is what lets the full latent, rather than a 3-D projection of it, be compared against physical 3-D. Place cells are those with significant Pearson correlation between rate maps (per-cell one-sided *P <* 0.025 against 100 circular-shift shuffles of held-out flights, peak rate *≥* 1 Hz, and finite correlation); azimuth- and goal-heading-tuned cells use the identicalcriterion on the corresponding 24-bin heading ring. Permutation tests use 2,000 iterations; the PV-vs.lag bands use 500 place-cell bootstraps. See also Figs. S8–S9 for additional analysis and examples.

**Fig. 4.**
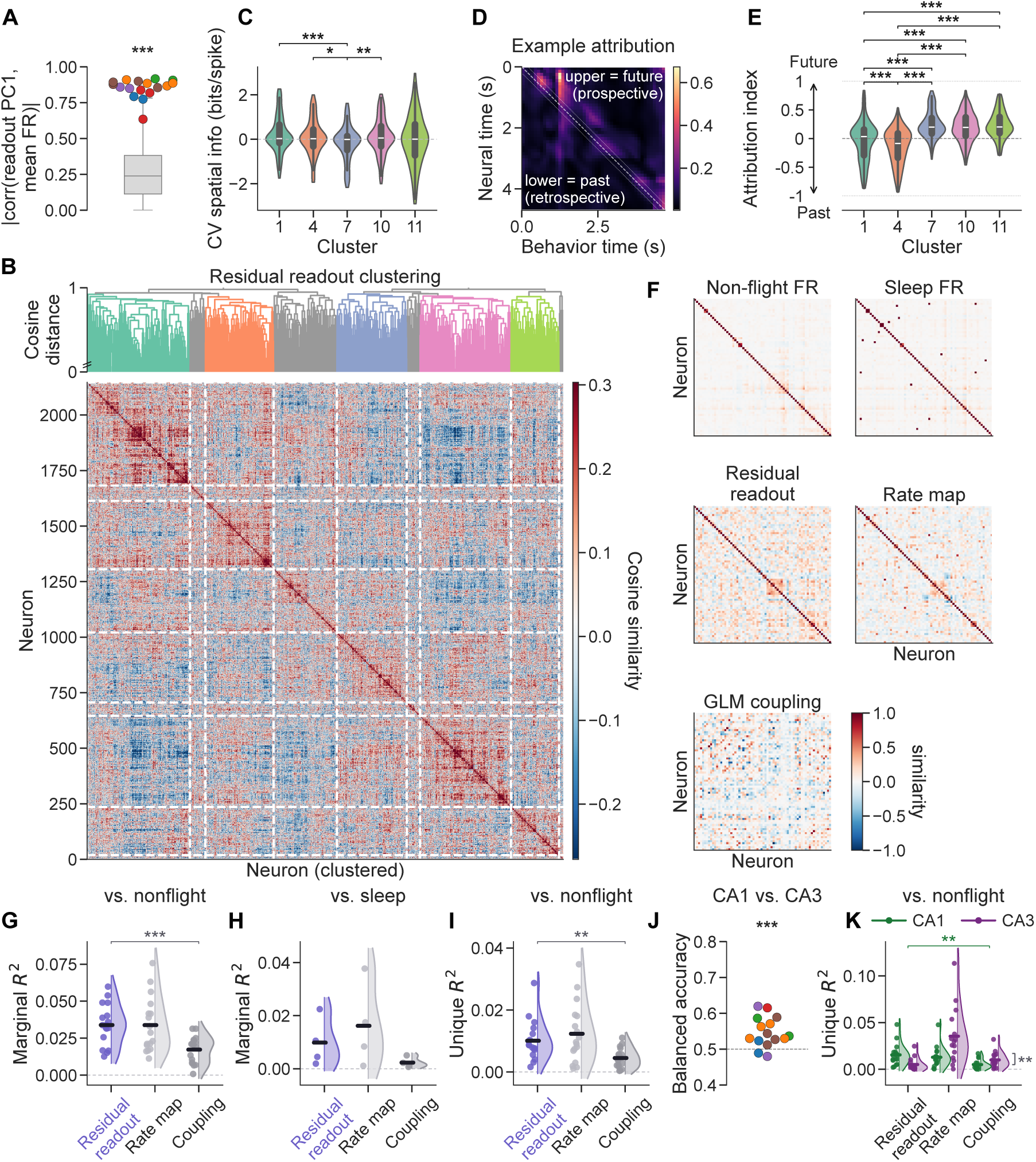
Neuronal readout weights encapsulate known and novel biological structure. (A) The absolute Pearson correlation between the first principal component (PC1) of each session’s readout weight matrix and that session’s per-neuron mean firing rate (FR; one point per session, colored by bat; *n* = 17 sessions). Gray box = random-direction null per session; two-sided Wilcoxon signed-rank test comparing the observed per-session absolute correlation to its null median (*P* = 1.53*×*10*^−^*^5^). (B) Hierarchical clustering (average linkage, cosine distance) of the residual readout weights (each neuron’s unit-normed readout vector orthogonalized against the population mean-firing-rate axis) across the 2,147 well-fit neurons (Poisson FDE *>* 0). Dendrogram (*top*) and neuron*×*neuron cosine-similarity matrix (*bottom*), sharing the cluster-ordered neuron axis; dashed white lines mark the boundaries between the five well-populated clusters (colored). (C) Cross-validated spatial information (bits per held-out spike) per neuron, grouped by readout cluster. Kruskal–Wallis test across clusters overall (*P* = 7.06 *×* 10*^−^*^5^); brackets show Holm-corrected pairwise two-sided Mann–Whitney U tests (*n* = 183–419 neurons per cluster). Cluster 7 carries less cross-validated spatial information than clusters 1 (*P* = 1.05 *×* 10*^−^*^5^), 10 (*P* = 6.7 *×* 10*^−^*^3^) and 4 (*P* = 0.037); no other pair differs (1 vs. 11 *P* = 0.30;1 vs. 4 and 7 vs. 11 *P* = 0.37; 1 vs. 10 *P* = 0.58; 4 vs. 10, 4 vs. 11 and 10 vs. 11 *P* = 1.00). (D) Example neural*→*position attribution (Jacobian) matrix over time for one test flight; upper triangle = future position (prospective), lower = past (retrospective). Color = ∥*∂* rate*/∂* position ∥_2_; dashed line = neural time equals behavior time; dotted lines = *±*200 ms exclusion band around the diagonal. (E) Future-past attribution index (pooled across test flights; *>* 0 = prospective) per neuron, grouped by readout cluster; dashed line at 0; *±*1 marked. Brackets are Holm-corrected pairwise two-sided Mann–Whitney U tests between readout clusters (*n* = 217–464 neurons per cluster): clusters 1 and 4 each differ significantly from clusters 7, 10 and 11 and from one another (1 vs. 4 *P* = 7.1 *×* 10*^−^*^5^; 4 vs. 7 *P <* 10*^−^*^300^, 4 vs. 10 *P* = 3.0 *×* 10*^−^*^40^, 4 vs. 11 *P* = 3.5 *×* 10*^−^*^36^; 1 vs. 7 *P* = 2.3 *×* 10*^−^*^30^, 1 vs. 10 *P* = 9.6 *×* 10*^−^*^23^, 1 vs. 11 *P* = 3.4 *×* 10*^−^*^21^), whereas clusters 7, 10 and 11 do not differ (7 vs. 10 *P* = 0.054; 10 vs. 11 and 7 vs. 11 *P* = 0.48). (F) Example sleep-matched session: non-flight FR, sleep FR, residual-readout, rate-map and GLM-coupling similarity matrices over matched neurons, sorted by the non-flight rest clustering order on a common color scale. (G) Multiple regression on similarity matrices (MRM) against non-flight rest noise correlations: marginal *R*^2^ of each predictor (residual readout, rate map, coupling) entered alone. Brackets: paired session-level two-sided Wilcoxon signed-rank tests of each competitor vs. the residual readout reference (residual readout vs. rate map *P* = 0.611, n.s.; vs. coupling *P* = 1.53 *×* 10*^−^*^5^; *n* = 17 sessions). (H) The same marginal *R*^2^, but with the regressand swapped from non-flight rest to sleep noise correlations, over the *n* = 4 sessions with a matched sleep recording (205 of 487 well-fit neurons matched across the two recordings; 3 bats). (I) MRM unique (semipartial) *R*^2^ of each predictor against non-flight rest, holding the other two fixed (residual readout vs. rate map *P* = 0.284, n.s.; vs. coupling *P* = 1.34 *×* 10*^−^*^3^; *n* = 17 sessions). (J) A per-session RBF SVC classifies CA1 vs. CA3 membership from residual readouts; one balanced-accuracy point per session (colored by bat) across the *n* = 15 sessions with at least 10 neurons per region, session-level two-sided Wilcoxon signed-rank test vs. chance = 0.5 (*P* = 4.27 *×* 10*^−^*^4^). (K) MRM unique (semipartial) *R*^2^ of each predictor computed separately within CA1 (green) and CA3 (purple); points = individual sessions, half-violins their distribution, thick bars the across-session mean. Green brackets: the CA1 simple effect (residual readout *>* competitors), paired session-level two-sided Wilcoxon signed-rank tests (residual readout vs. rate map *P* = 0.548, n.s.; vs. coupling *P* = 9.34 *×* 10*^−^*^3^; *n* = 17 sessions with sufficient CA1 neurons). Black asterisks beside the vertical bracket connecting the CA1 and CA3 means mark the two-way repeated-measures ANOVA predictor *×* subregion interaction (Greenhouse–Geisser corrected *F*_2,28_ = 9.47, *P* = 2.79 *×* 10*^−^*^3^; *n* = 15 sessions present in both regions). See also Fig. S10.

Our model outperformed many alternatives at both decoding and encoding, including both traditional and deep network-based approaches (Fig. S1D–H; [62]). Decoded flight trajectories closely followed the observed three-dimensional flight paths (Fig. 1B), substantially outperforming a näıve Bayes decoder [63] and modestly but significantly outperforming a CEBRA-based decoder [64], as quantified by Fŕechet distance (Fig. 1C). For neural encoding, the model reproduced trajectory-specific firing-rate patterns across diverse flight paths (Fig. 1D). Following previous observations that place coding in flying bats is trajectory-specific [42,45], we clustered flights according to their empirical Fŕechet distance and constructed both place-based and trajectory-specific generalized linear models (GLMs), with the latter testing the hypothesis that spatial coding is trajectory-specific but otherwise static and linear [65]. Across sessions, the Twin outperformed both variants of GLMs, as measured by Poisson fraction deviance explained (FDE) and Pearson correlation (Fig. 1E–F).

Even when augmented with fixed time lags, the GLMs continued to fail to match the Twin’s performance (Fig. S1E–F). Adding neuronal history and coupling filters [66] restored comparable encoding performance but substantially changed the information available to the models by allowing them to exploit autocorrelations and correlations among simultaneously recorded neurons (Fig. S1G– H). Surrogate analyses indicated that these correlation-aware models relied more heavily on spike correlations, whereas the Twin was more sensitive to behaviorally structured patterns extending across the flight (Fig. S2). Consistent with this interpretation, restricting the model’s temporal con-text window during training impaired both encoding and decoding performance relative to training on complete flights (Fig. S3A). Echoing previous neural foundation models [18–21], we found that our architecture transferred well to held-out sessions; fine-tuning only the session-specific readout weights while freezing the core outperformed training on new data from scratch and approached performance of the full model (Fig. 1G–H). Conversely, reducing the total number of training sessions impaired encoding performance (Fig. S3B).

Due to its predictive accuracy and joint architecture in which neural and behavioral tokens are embedded and concatenated, we refer to our model throughout the manuscript as a functional digital “Twin”. This terminology is not meant to imply that the model exhibits all the same characteristics of the mammalian hippocampus, nor that it perfectly predicts held-out data. Rather, we use the term to emphasize the experimental accessibility of the Twin as a means of generating testable hypotheses about the organizing principles of hippocampal activity, rather than simply as a tool for statistical prediction.

### 3.2 Population activity emphasizes flight motif over position

Hippocampal place representations [32,33] are well-known to deviate from strict Euclidean geometry across species [34–48]. Historically, scientists have identified such deviations by testing specific hypotheses through carefully designed experiments. We asked whether the Twin’s learned latent space could reveal the organization of hippocampal population activity during natural behavior without specifying the relevant organizing variables in advance. We therefore examined how three-dimensional flight trajectories were represented in this space, asking whether spatial dimensions were expanded or compressed, trajectories were nonlinearly reshaped, or position was represented jointly with other behavioral variables (Fig. 2A-E).

The digital Twin learns a latent representation from which three-dimensional position is decoded linearly and individual neuronal firing rates are predicted through a linear readout followed by a fixed nonlinearity. This simple readout allows us to examine how the learned representation organizes behavior to account for neuronal responses. We compared this representation with CEBRA [64], a nonlinear contrastive learning method increasingly used to study behaviorally relevant structure in neural activity [67,68]. Both methods used three-dimensional position labels and underwent similar hyperparameter optimization, but the labels served different roles. In the CEBRA configuration used here, they guided the selection of observations brought together through contrastive learning. In the Twin, position served as an input for predicting neural activity and a target for behavioral decoding. These approaches offer complementary perspectives on neural organization: one emphasizes relationships defined by selected behavioral variables, while the other learns features that support direct prediction of neural activity and behavior.

Both CEBRA and the Twin represented flight trajectories along curved structures in latent space (Fig. 2F), but differed in how they organized flights relative to one another. Whereas CEBRA mapped identical physical locations to proximal points on the manifold, the Twin organized flights into two distinct lobes (Fig. 2G). The two lobes corresponded to flight segments traveling in oppo-site directions along the principal flight axis, which closely aligned with the longest axis of the flight enclosures (Fig. S4A-D), and which we call flight “motifs.” Even long, looping flights that shared an origin and destination began on the Twin’s first lobe before switching to the second, whereas no such structure was apparent in the CEBRA embedding (Fig. 2G, Fig. S4E). Neural-token representations derived from behavior during encoding and from recorded spikes during decoding occupied overlapping regions of latent space, with the behavior-derived trajectories appearing smoother but otherwise exhibiting similar structure (Fig. 2G, Fig. S5).

Within each lobe, spatially distinct flight segments followed similar latent trajectories, suggesting recurring patterns of population activity. These observations raised questions about how the shared organization relates to reported goal-vector ([41], but see [42,69]), reward-relative [60], and direction-dependent place coding [34–36,43,47]. The organization was evident during flights toward both reward sites and perches, indicating that it was not restricted to reward approach (Fig. S4E). Direction-dependent place coding could account for the separation of outbound and return flights, but would not by itself explain the alignment of spatially distinct paths within each lobe (Fig. 2C).

These observations led us to hypothesize that the recurring structure in the Twin’s latent space reflects a shared organizing principle of hippocampal activity across different flight paths. We next quantified whether flight segments traveling in the same direction occupied similar regions of the full-dimensional latent space across different flight paths and animals, and whether this shared organization could be observed at the level of single-neuron tuning and neural population sequences.

### 3.3 Latent representations align neural geometry and dynamics across animals

To test this prediction emerging from analysis of the digital Twin, we turned to representational similarity analysis (RSA). We computed the symmetric Hausdorff distance between the 3-D (behavioral) or 64-D (latent) vectors for each indoor flight segment. This metric accommodates flights of different lengths and is minimized when every point of either set lies close to a point in the other set. We then z-scored the distances within each representational space and arranged them into symmetric matrices. Because these segments spatially overlap, RSA in the physical space failed to reveal significant structure (Fig. 2F–H, *left column*). CEBRA picked up on structure specific to particular routes, bats, or flight clusters, but failed to cleanly distinguish motifs (Fig. 2F–H, *second column*). By contrast, RSA on our Twin exhibited block-diagonal structure at the motif level which generalized across animals and routes (Fig. 2F–H, *right two columns*). Consistent with this organization, four landmark positions — corresponding to flight origins and destinations at the reward and the perch — while entangled in the physical space, mapped to four linearly separable quadrants in the Twin’s latent space (Fig. S6A–E). Note that this motif structure with two lobes as well as the decodable landmark positions in the latent space were a byproduct of the Twin jointly modeling hippocampal activity and flight kinematics, emerging spontaneously without an explicit optimization objective.

Comparing the Hausdorff distance within versus across motifs showed motif separation was substantially stronger in the Twin latent space than in either physical space or the CEBRA latent space (Fig. 2I). Likewise, motif could be decoded from the time-averaged Twin embeddings using a leave-one-route-out analysis, substantially outperforming both the CEBRA embedding and animal-specific raw neural activity (Fig. 2J). Other continuous kinematic variables could also be decoded from the Twin representation (Fig. S6F). Thus, our functional digital Twin was able to extract shared latent structure that generalized across animals and routes, both in the lab and in an out-door field station, and supported downstream decoding, beyond what was possible using either the untransformed neural activity or existing baselines.

In addition to the geometry of neural latent representations, the Twin allowed us to analyze their dynamics on a segment-by-segment basis. To extract temporal and spatial patterns, we performed dynamic mode decomposition (DMD; Fig. 2K), a data-driven technique originally developed to identify coherent dynamical features from complex fluid systems and more recently applied to neural population dynamics [70–73]. DMD decomposes time-series data into dynamic modes, representing recruited neural ensembles, and their associated eigenvalues, which characterize their continuous temporal evolution, including frequency and decay or growth rates (see Methods). We observed that most estimated eigenvalues were located within or near the boundary of the unit circle in the complex plane, indicating stable, non-divergent population dynamics (Fig. S7). Furthermore, the power spectrum of these modes exhibited characteristic scaling, with negative spectral slopes steeper than *−*1.0 (Fig. S7), consistent with stable, long-range temporal correlations in the latent space.

To compare motif-level dynamics across trajectories, we constructed temporal dynamic profiles by binning the complex plane, weighting each bin by the corresponding mode power, and flattening the resulting distributions. For spatial alignment analyses across motifs, we evaluated the spatial profile (the real component) of the highest-power oscillatory DMD mode. Interestingly, temporal profiles were consistently similar across flight segments (Fig. 2L), indicating shared temporal dynamics across motifs. By contrast, the similarity matrix for spatial DMD modes displayed a distinct block structure: spatial profiles were highly correlated within the same motif but remained largely orthogonal across motifs (Fig. 2M), with motif identity represented substantially more strongly in the spatial than the temporal profiles (Fig. 2N). Thus, motif identity was expressed primarily through the spatial organization of the Twin’s latent dynamics, whereas their temporal evolution was largely shared across trajectories.

### 3.4 Latent representations expose spatial generalization at the single-neuron and population level

Previous studies have demonstrated that bat hippocampal neurons maintain highly stable spatial firing across days, but that this activity pattern depends on the specific flight pattern executed by the bat [42]. Thus, whether and how neural tuning is related across *different* flight patterns remains unclear. Our population analyses revealed that flights sharing the same motifs preserved similar latent geometry and dynamics, prompting us to examine the corresponding organization at the single-neuron and population-levels.

We reasoned that if our manifold view captures a core aspect of bat hippocampal single-neuron tuning, then place cell sequences should be preserved across *different* flight paths sharing the *same motif(s)*, even when their spatial positions and directions differ substantially. Note that this is the converse of the typical analysis, wherein the instantaneous *spatial position*, and sometimes direction, is held fixed, while the *overall path* is allowed to differ. In this latter case, different routes to the same goal, or different directions of travel entirely, are generally believed to result in at least partial “remapping”, yielding orthogonal place cell representations, including along shared positional segments [34–39,44,48]. By contrast, generalization across spatially distinct paths sharing the same motifs would be counterintuitive, although not ruled out by existing data (cf. [40]).

We therefore identified flight clusters with similar origins and destinations but markedly different flight paths and directions (Fig. 3A). Consistent with the prediction emerging from the digital Twin, neurons that fired strongly at a particular position along Path 1 could be silent at that exact position on Path 2, instead forming a new, spatially distant firing field (Fig. 3B). Importantly, this reorganization was not random: tuning was preserved as a function of normalized flight progress (Fig. 3C). Across the population, place cells, identified by significant held-out correlation on the Path 1 flight cluster (see Methods), formed a repeatable sequence not only along Path 1 (Fig. 3D, *left*), but also along multiple other paths sharing the same motifs (Fig. 3D, *right*). Computing the population vector (PV) correlation across paths at each combination of lags revealed a clear concentration along the main diagonal at zero lag (Fig. 3E), which could not be explained by Euclidean distance (Fig. 3F) or goal heading vector (Fig. 3G) coding. In other words, hippocampal population activity evolved similarly as the bat proceeded along each path, despite the paths being spatially and directionally distinct. Consistent with this, PV correlation tracked normalized flight progress better than either azimuth or goal heading (Fig. 3H). Likewise, peak-progress ordering across place cells was significantly preserved between paths, whereas peak ordering by azimuth or goal heading was not (Fig. 3I, Fig. S8). Additional flight cluster pairs from other sessions exhibited the same overall trends (Fig. S9).

In the digital Twin (Fig. 3J), the same example unit’s diffuse physical place fields (Fig. 3B) converged into a compact region of the latent space (see more examples in Fig. S9). Based on this observation, we systematically quantified the spatial similarity of neuronal place fields between flight-cluster pairs with the same route identity, comparing the three-dimensional physical space against the space spanned by the first three principal components (PCs) of the neural latent representations. Treating each rate map as a spatial distribution, we evaluated cross-cluster similarity using the 3-D Wasserstein distance (see Methods). Higher values indicate greater distance of firing fields across flight clusters with the same route identity. The majority of neuron-route pairs exhibited lower Wasserstein distances in the latent space compared to the physical space, demonstrating that our Twin learns to collocate physically distant firing fields within a compact region of latent space (Fig. 3K).

Lastly, to evaluate neural tuning fidelity across routes, animals, and environments, we calculated the difference in these similarity metrics between the latent and physical spaces. In all cases, this difference was significantly different from zero (Fig. 3L), indicating that single-neuron firing fields were more consistently aligned across flight clusters belonging to the same route in latent space than physical space. The similarity measured in latent space also generally exceeded that obtained from the shuffled data (Fig. 3L,M, Methods). Together, these results suggest that hippocampal single-neuron tuning across spatially distinct trajectories is better preserved in the population-derived latent representation than in raw physical coordinate space.

### 3.5 Neuron-specific readouts reveal functional organization across brain states

The preceding sections establish the utility of the digital Twin’s latent embeddings for characterizing neuronal activity across flights as a function of the behavioral inputs. We next asked whether the Twin’s parameters, and specifically the session-specific neuronal readouts that remain fixed across flights, capture stable functional properties of the recorded neurons. Although these parameters are not direct proxies for biological synaptic weights, they map the model’s latent activations onto the activity of individual recorded neurons through a linear transformation (with a static nonlinearity) and may therefore contain information about each cell’s anatomical location or functional role [21].

We first observed that the first principal component (PC1) of the readout weights correlated strongly with each neuron’s mean firing rate, likely because the neuron-specific bias term could capture a baseline offset but not the firing rate modulation that scaled with it (Fig. 4A). Because this dominant component largely reflected mean firing rate rather than more specific neuronal properties, we restricted the analysis to well-predicted neurons (Poisson FDE *>* 0), orthogonalized each neuron’s readout vector with respect to the population mean firing rate axis, and hierarchically clustered the resulting “residual readout” matrix (Fig. 4B). The residual readouts exhibited substantial structure relative to appropriate controls (Fig. S10A–B), revealing functional clusters that were conserved across sessions and animals (Fig. S10C) while differing in their cross-validated spatial information (Fig. 4C), encoding performance, average firing rate, and dorsoventral position (Fig. S10D–I).

More compellingly, these functional clusters captured aspects of neuronal tuning that were never explicitly labeled or used as training targets. For each test flight, we computed the Jacobian of every neuron’s predicted rate with respect to the bat’s spatial position (Fig. 4D) and then compared the degree of retrospective (lower triangle) to prospective (upper triangle) coding (“attribution index”, see Methods). We found that clusters differed on average in their relative weighting of past and future (Fig. 4E), suggesting that temporal context is integrated within the latent space and then differentially read out by distinct groups of neurons on a moment-by-moment basis, consistent with previous findings from the hippocampus of freely flying bats [74].

The residual readout matrix also contained information that did not depend on functional clusters. For each simultaneously recorded neural population, we compared the cosine-similarity matrix of its residual readouts with several measures of pairwise neuronal similarity (Fig. 4F). We computed the Pearson correlation between neuronal spike counts during non-flight epochs — either waking rest within the session, or, for a subset of sessions, sleep preceding the session. We then compared how well residual-readout similarity, spatial rate-map correlation, and the time-integrated, symmetrized cross-neuron coupling weights from a conventional GLM predicted neuronal correlations across these distinct brain states (see Methods). Note that neural activity from these non-flight and sleep epochs was not used to train the digital Twin or any of the other predictors.

To quantify these relationships, we performed multiple regression on similarity matrices (MRM) and compared the variance explained by each predictor, both alone and after controlling for the others. Specifically, we vectorized the upper triangle of the non-flight firing rate correlation matrix and those of the residual-readout, rate map, and GLM-coupling similarity matrices, rank-standardized each entry, and regressed the target vector on the predictor vector or vectors. Residual-readout similarity explained more variance in non-flight firing-rate correlations than GLM coupling strengths, and a similar amount as rate-map similarity (Fig. 4G–H). The utility of the residual readouts remained even when considering only the unique variance contributed by each predictor (Fig. 4I). Thus, functional relationships learned by the Twin exclusively from flight activity generalized to neuronal correlation structure during waking rest and sleep, on timescales of hundreds of milliseconds, over and above what is obtainable from baseline methods such as rate maps.

Finally, we analyzed CA1 and CA3 separately, motivated by our ability to classify each neuron’s hippocampal subregion above chance from its residual readout (Fig. 4J). Because CA1 activity is strongly shaped by afferent inputs, we hypothesized that residual-readout similarity, which reflects how individual neurons map shared latent input onto firing rates without explicit cross-neuron coupling, would better capture its correlation structure. By contrast, the dense recurrent connectivity of CA3 should be more directly reflected in explicit cross-neuron coupling terms [75]. Consistent with this hypothesis, residual-readout similarity accounted for more unique variance than did coupling strength in CA1, and the subregion *×* predictor interaction was significant (Fig. 4K). These results support the conclusion that the digital Twin and coupled GLMs capture complementary sources of neuronal covariance, with CA1 benefiting particularly from the Twin’s shared latent representation (Figs. S1–S2).

## 4 Discussion

### 4.1 Discovering latent structure during natural behavior with a digital Twin

Recent influential models of the hippocampus emphasize relational and sequential structure, with physical space viewed as one manifestation of a broader representational framework [49,50,76–78]. This perspective can account for place or grid-cell-like activity across a wide range of non-spatial domains [51–60], as well as systematic deviations from purely Euclidean spatial coding [34–48].

A central challenge, however, is to discover from neural data the particular latent structure that animals deploy in a given environment. This challenge is especially acute during natural behavior, where the experimenter does not define the task structure and may not know in advance which features of behavior organize the underlying neural activity [1,79]. Recent foundation-model and digital-twin approaches have demonstrated strong predictive performance and generalization across animals and stimulus domains [21], but have focused primarily on sensory responses in controlled, head-fixed paradigms. Whether these models can generate testable hypotheses about higher-order cognitive function during self-directed natural behavior has remained unclear. Here we demonstrate this possibility in the hippocampus.

Our functional digital Twin provides a scalable, high-fidelity method for extracting latent hippocampal structure across sessions, animals, and environments. Interrogating its internal representations revealed that hippocampal activity was organized around recurring flight motifs and generated the unexpected prediction that neural sequences should be preserved across spatially distinct trajectories. Analysis of its neuron-specific readouts further identified functional relationships that generalized beyond flight to waking rest and sleep — brain states not used to train the model. We then tested and validated these model-derived predictions directly in the recorded neural activity. Together, these findings demonstrate that predictive models can serve not only as accurate descriptions of brain–behavior relationships, but also as experimentally accessible tools for generating and testing biological hypotheses during natural behavior.

### 4.2 Latent structure across space, time, and internal states

Interrogating the digital Twin reveals a form of hippocampal organization that reconciles context sensitivity with generalization. Consistent with previous reports of splitter-like coding [34–39,48], neuronal activity at a given physical location depended on direction of travel and temporal context. Yet, the *same* hippocampal neuron could also exhibit corresponding tuning at spatially *distinct* locations when those locations occupied similar positions within a shared flight motif in the latent space. At the population level, the ordering of hippocampal sequences was preserved across trajectories that differed markedly in physical position and heading. Thus, apparent remapping in physical space can coexist with conserved organization in an underlying sequence space. This organization is reminiscent of representations of progress toward a goal or action plan [46,47], but here emerged during self-directed, three-dimensional navigation. In contrast to classical time-cell coding [51,80–82], the relevant coordinate here was the progression along an ongoing self-directed trajectory rather than elapsed time, and the sequence structure generalized across spatially distinct trajectories sharing a recurring behavioral motif but not across distinct motifs.

At first glance, this generalization may seem inconsistent with rodent studies showing that hip-pocampal representations can be specific not only to a combination of origin and destination, but also to the complete route traveled between them. However, such route specificity may depend strongly on experimenter-imposed task demands, including differential reward across routes [40,44]. Our results suggest that when animals select routes freely, without one route being explicitly favored over another, route-specific spatial firing can coexist with a shared sequential organization, consistent with broader generalization [50]. More generally, the latent structure of hippocampal representations is likely to vary across species, tasks and environments according to the navigational strategies animals deploy. The digital-twin approach provides a data-driven means of discovering this structure, without requiring the experimenter to specify in advance whether the relevant coordinate is position, direction, goal, elapsed time or another behavioral variable. Applied across contexts, this strategy may reveal both shared and species-specific organizing principles underlying the diversity of functional response types observed throughout the hippocampal formation [49,50].

The digital Twin also exposed functional structure that generalized across brain states. Although the model was trained exclusively on activity during flight, similarity between neuron-specific residual readouts predicted pairwise neuronal correlations during both waking rest and sleep more effectively than the coupling weights of conventional generalized linear models [66], adding information over and above rate-map similarity. This relationship was particularly strong in CA1, whereas explicit cross-neuron coupling better captured correlation structure in recurrently connected CA3, consistent with the distinct circuit organization of the two subregions [75]. These findings suggest that the Twin’s readouts capture stable, input-related aspects of neuronal organization that are expressed across behavior and internal states, whereas coupled GLMs preferentially capture recurrent interactions among neurons.

Importantly, the identification of these latent structures was robust to variation in both model architecture and experimental data. Consistent functional manifolds were observed across multiple recording sessions and across distinct computational approaches, including pure encoding frameworks predicting neural responses directly from bat behavior. These results demonstrate that the emergent functional structure is a fundamental property of well-trained predictive models rather than an artifact of a specific architecture or session.

### 4.3 Representational correspondence without algorithmic equivalence

A substantial body of work has shown that artificial neural networks trained to perform path integration can develop units with place-, grid-, and other spatially tuned responses [83], including both recurrent neural networks (RNNs) [84,85] and transformers [86]. Transformers are particularly interesting in this context because the attention mechanism has a formal correspondence to the update rule of modern Hopfield networks, which themselves are closely related to longstanding theoretical accounts of hippocampal associative memory [87,88].

These formal correspondences do not imply that the hippocampus implements transformer attention algorithmically or that our digital Twin reproduces its biological mechanisms. Nor do we establish that the model’s performance relative to alternative architectures, including RNNs, arises from the relationship between attention and associative memory. A more parsimonious explanation is that the general strength of transformers in sequence modeling [61] extends to integrating neural and behavioral information over long timescales [15–21]. At a functional level, however, this capability aligns with theoretical accounts in which the hippocampus constructs and relates sequences of latent states [49,50,77,78]. This shared computational demand may partially explain why the trans-former’s internal representations proved useful for revealing sequential structure in hippocampal activity, from which we could derive biologically testable hypotheses.

### 4.4 Scientific discovery via interpretability of digital Twins

Using AI to advance science has long been a goal across scientific disciplines [89–91], with notable successes in fields such as genomics [92] and protein structure prediction [11]. Yet increasing predictive power has not necessarily made it easier to extract general scientific principles from these models. Indeed, as datasets and model complexity grow, the knowledge captured during training can become progressively more difficult to extract. The emerging field of neural-network interpretability [93] seeks to reverse engineer the representations and computational mechanisms learned by artificial models, particularly in vision [94] and language [95,96]. For scientific models, however, interpreting the artificial network is not the endpoint. The central challenge is to translate what the model has learned into testable hypotheses about the natural system it represents.

This form of scientific interpretability requires fluency in both machine learning and the relevant biological domain. Some of the structure learned by a model may reflect its architecture, variables missing from the dataset, or incidental correlations rather than the organization of the biological system itself. Inferences from a digital twin should therefore be regarded as provisional until tested against biological data, either through independent post-hoc analyses, as we have done here, or closed-loop experimental designs. Importantly, digital twins can serve as *in silico* testing grounds to synthesize optimal perturbation protocols, such as most exciting inputs, to drive target neurons in real time [13,22–24]. Digital twins are thus not replacements for direct observation or perturbation. Neither are multimodal, multitask models wholesale replacements for smaller, simpler, intrinsically interpretable computational frameworks [8]. The former can capture the complexity and diversity of natural behavior and help identify candidate organizing structure, whereas the latter remain essential for isolating mechanisms and developing formal theory. Used together, these approaches offer a path from large-scale prediction and hypothesis generation to mechanistic explanation to *in vivo* perturbation and confirmation.

One salient advantage in the present context is the dimensionality of the behavior in question. Our Twin model operates on a 64-dimensional feature vector representing spatial-behavioral state, for which the first three principal components capture a sizable fraction of total variance. Digital twins of biological vision [21], by contrast, need to navigate multi-dimensional latent tensors — spanning spatial height, width, temporal depth, and feature channels — making latent space structure difficult to understand. In this way, the putative complexity of unconstrained spatial behavior in fact becomes a crucial simplification which allowed us to make progress directly through dimensionality reduction and visualization.

### 4.5 Limitations and future directions

We leveraged a unique dataset of multi-probe Neuropixels recordings in freely flying bats, comprising thousands of hippocampal neurons recorded across indoor and outdoor environments [45]. Although the Twin outperformed the alternative models we tested on both neural encoding and behavioral decoding, it remains necessarily incomplete. Because the same neurons were not recorded across environments, we could not determine how individual neuronal representations remapped between them. Moreover, the model received only hippocampal activity and three-dimensional position and therefore lacked many variables that could influence hippocampal dynamics during natural flight, including head orientation, sensory and echolocation signals, motor state, reward expectation and other internal variables. Consistent with this incomplete access, a substantial fraction of neural variability remained unexplained. We therefore use “digital twin” in a functional and deliberately limited sense: the model captures reproducible relationships between measured neural activity and behavior but does not constitute a complete replica of hippocampal circuitry or computation.

Data scale presents an additional limitation. Even recent models trained on millions of neurons recorded from dozens of head-fixed mice — amounting to approximately 150 billion neural tokens — remain limited by the quantity of available data and fall well short of perfect prediction [20]. Although our model demonstrates transfer to held-out recording sessions (Fig. 1G–H), substantially larger datasets will be needed to improve predictive fidelity, increase model capacity and determine how broadly the discovered representations generalize [97–99]. Longitudinal recordings of the same neurons across environments and behaviors would be particularly valuable, allowing the direct test of whether the latent structure identified here predicts remapping, generalization and the emergence of new representations.

At the same time, neuroscience models must scale not only in the quantity of data, parameters and computation [100], but also in the diversity of the behaviors and biological contexts represented. Computations expressed specifically during self-directed three-dimensional navigation cannot be recovered from datasets in which such behaviors never occur, regardless of their scale. Natural behavior therefore represents an additional and particularly important axis of scaling for biological foundation models. On the data side, richer behavioral measurements, longer recordings, additional brain regions, new environments and a broader range of species should both improve predictive performance and expand the range of biological hypotheses that digital twins can generate. On the model side, semi-supervised objectives, better multimodal and multitask training, improved handling of missing inputs and outputs, and — more speculatively — embodiment will allow these approaches to integrate datasets that can differ more widely in their measured behavioral variables or neural recording techniques [101,102]. Together, these advances may ultimately produce qualitative transitions in model capabilities [103,104], enabling digital twins to progress from retrospective analysis of existing datasets towards prospective prediction and experimental design.

## 5 Methods

### 5.1 Neural data collection

All data analyzed in this study were previously collected and reported [45]. Briefly, six adult male Egyptian fruit bats (*Rousettus aegyptiacus*) were anesthetized, mounted on a stereotaxic apparatus (Model 942; Kopf Instruments) and implanted with a training cone. During the same surgery, craniotomies were drilled targeting dorsal hippocampus (6.3 mm anterior to the transverse sinus that runs between the posterior part of the cortex and the cerebellum and 3.2 mm lateral to the midline). Seven to ten days later, 2-3 Neuropixels 1.0 probes were coated with fluorescent dye (CM -DiI; Invitrogen C7001), inserted into the craniotomies to a depth of 5500 *µ*m, and cemented in place using dental acrylic. Recordings began one day after probe insertion and were performed using a SpikeGadgets wireless Neuropixels 1.0 headstage. Drift correction and spike sorting was done automatically offline using Kilosort4 [105]. At the end of the experiment, bats were given a lethal overdose of sodium pentobarbital and transcardially perfused with 3.7% paraformaldehyde. The brain was removed, sectioned, and imaged using an Axioscan Slide Scanner (Zeiss) to localize the probe track. All probes were successfully identified in the hippocampus. All animal protocols were approved by the Animal Care and Use Committee (ACUC) of the University of California, Berkeley.

### 5.2 Aerial foraging

Foraging experiments took place in an indoor flight room (*n* = 4 bats, 5.6 m *×* 5.2 m *×* 2.5 m) or an outdoor flight enclosure (*n* = 2 bats, 10.1 m *×* 4.1 m *×* 2.8 m). Indoors, the 3D spatial position of the bats was tracked at millimeter resolution at 120 Hz using 16 motion-capture cameras (Raptor-12HS, Motion Analysis). Outdoors, the 3D spatial position of each bat was recorded at 100 Hz using an ultra-wideband based real-time location system (Ciholas). Two feeders indoors (or one feeder outdoors) were placed along one wall and dispensed puŕeed fruit reward when a bat landed on the feeding platform, controlled either by an infrared beam break sensor (indoors) or remotely by the experimenter (outdoors) via an Arduino (Uno Rev3) and Adafruit Motorshield (1438; Adafruit).

### 5.3 Sleep recordings

Sleep recordings were obtained from a subset of animals (3 bats, 4 sessions). Bats were housed under a reversed light cycle (lights off from 07:00 to 19:00), with active behavior occurring during the dark phase and sleep during the light phase. Sleep recording sessions of *∼*1h duration were performed approximately 2 h after the onset of the light phase (from *∼*21:00). Each implanted bat was placed individually in a small transparent cage inside a sound-attenuated room. Animals were continuously monitored with multiple cameras for behavioral signs of sleep or quiescence, including eyelid closure and the absence of gross body movements. Neural data and accelerometer signals were acquired using the same recording hardware and software as during awake sessions. We confirmed that a large fraction of each session involved sleep based on converging evidence from video monitoring, absence of movement in the accelerometer signal, and presence of slow oscillations in the LFP, and so we did not further segment sleep sessions.

### 5.4 Data preprocessing

Spikes were counted and positions were averaged within 100 ms bins across the entire session. Flights were identified on the basis of a velocity threshold of 0.5 m/s before binning, and bins that included flight start and end were included as flight bins. Analyses were restricted to flight bins except where noted. Behavioral channels were *z*-scored per session using training-split statistics. Each flight was one sample, zero-padded to the longest flight of its session and carrying a validity mask, and all losses and metrics were evaluated on valid timesteps only.

Individual flights were resampled to seven points, compared by discrete Fŕechet distance and grouped into trajectory clusters by single-linkage clustering, cut at 0.8 indoors and 1.2 outdoors, following [45]. Flights whose peak acceleration exceeded 50 m s*^−^*^2^ were discarded as tracking artifacts. Within each session, flights were split 70/15/15 into training, validation and test sets with a fixed seed; this split is shared by every model and analysis in the paper, and the test split was used only for reported results, never for fitting or model selection.

### 5.5 Digital twin architecture

Each 100 ms bin produced two tokens: a neural token from a per-session linear read-in of spike counts, and a behavioral token from a per-environment linear read-in of 3-D position. Tokens carried a learned modality embedding, and tokens serving as prediction targets had their content replaced by a shared learned mask token while remaining in the sequence. The two streams were concatenated into a length-2*T* sequence and passed through four pre-norm transformer blocks with full attention and timestamp-aware rotary position embeddings, so a neural and a behavioral token from the same bin share rotary phase; padded timesteps were excluded from attention. Latent dimension was 64, with four heads, feed-forward width 64, dropout 0.2 and untransformed spike counts.

Six masking schemes were sampled during training: encode and decode, in which all valid tokens of one modality are targets; within-modality infilling, masking 50% of timesteps in one modality with the other fully visible; cross-modal infilling, masking the same 50% in both; and two forecasting variants that predict the second half of the neural sequence and differ only in whether behavior is visible [19,20,106]. Only encode, decode and cross-modal infilling contributed to the model used throughout the paper (Table S1).

### 5.6 Training and model selection

The loss was a Poisson negative log-likelihood on masked neural targets plus a mean-squared error on masked behavioral targets scaled by *λ*, over valid timesteps only. The model was optimized using AdamW [107], with sessions interleaved in a randomized round-robin order within each epoch. After the end of each epoch, validation decoding error (mean across sessions of the median 3-D error in meters) and validation encoding fit (median across neurons of Poisson FDE) were recorded to track the Pareto front over the two. The optimal checkpoint was selected based on the joint score defined in Table S1, with early stopping after 40 consecutive epochs without a non-dominated update. Architecture, optimization and scheme-weight hyperparameters were selected by 200 Optuna trials of 300 epochs each, and the winning configuration retrained for 1,000 epochs to give the single model used throughout. Every hyperparameter search reported in this paper used Optuna and was scored on the validation split.

For the transfer analysis (Fig. 1G-H), each fold trained a base model on the remaining 16 sessions for 500 epochs, froze it, and then optimized only the held-out session’s neural read-in and readout for up to 500 further epochs. The from-scratch control trained the same architecture on the held-out session alone for 1,000 epochs. Transfer models used one fixed hyperparameter setting — that of the full model — with no search. All models were trained on a single NVIDIA GeForce RTX 5090 GPU.

### 5.7 Metrics, plotting conventions, and statistics

The Poisson fraction deviance explained (FDE) was defined as 1 *− D*(*y, y*^)*/D*(*y, y*), where *D*(*y, µ*) = 2Σ*_t_*[*y_t_*(log *y_t_ −* log *µ_t_*) *−* (*y_t_ − µ_t_*)] and *y* denotes the neuron’s mean count over the same time bins. For each neuron, encoding fit was evaluated across all valid test bins in a session using both FDE and the Pearson correlation between the observed and the predicted counts. Decoding accuracy was evaluated per flight using the discrete Fŕechet distance in meters.

For raincloud plots throughout the paper, dots show individual sessions (per-session medians), the half-violin their distribution, and the thick ink bar the across-session mean, with the model of interest in purple. The half-violin is a Gaussian kernel density estimate (bw method = 0.55), peak-normalized per-condition. In log panels (e.g. Fig. 1C), the KDE runs in log10 space. Violin plots in Fig. 4C,E and Fig. S10 trim points beyond the Tukey whiskers for legibility while all tests use untrimmed values.

Unless stated otherwise the unit of analysis was the session: per-neuron or per-flight values were reduced to a session median before testing. All tests are nonparametric and two-sided unless noted. Analyses used Python (v3.13.14) with NumPy (v2.4.6), SciPy (v1.18.0), pandas (v2.2.3), scikit-learn (v1.6.1), PyTorch (v2.7.1), and Optuna (v4.8.0). All random number generators were seeded from a single seed throughout.

### 5.8 Decoding baselines

The näıve Bayes place decoder [63] was fit per session on training flights as smoothed 3-D rate maps with an occupancy prior on a grid spanning the room, decoding position as the argmax of the Poisson log-posterior at *τ* = 100 ms. One hyperparameter set (grid resolution, smoothing, minimum occupancy, prior) was shared across sessions and chosen by 20 Optuna trials minimizing mean median 3-D validation error. Its continuity variant [63] replaces the flat prior with a Gaussian centered on the previously decoded point whose width scales with a fitted local mean-speed map, resetting at flight boundaries. Trajectory-conditioned variants fit one decoder per cluster with *≥* 3 training flights plus a catch-all over all remaining flights, routed at decode time by the flight’s cluster.

The unified CEBRA [64] model was trained across all 17 sessions with neurons concatenated, using the time delta conditional, 6,000 iterations and random neuron- and timestep-masking, with position read out by *k*-nearest-neighbor regression (*k* = 36, cosine). Hyperparameters (num hidden units, output dimension, time offsets, model architecture, temperature, and learning rate) were chosen by 40 Optuna trials. Note that this procedure decisively favored higher output dimensions, requiring post-hoc (linear) dimensionality reduction for visualization. A standard multi-session CEBRA was tuned by an independent search. The LSTM-RNN [62] used a per-session linear read-in onto a shared recurrent core and head, tuned by 20 Optuna trials with the published configuration enqueued as the first trial.

GLM-inversion decoders freeze a trained encoder and optimize the trajectory to maximize Poisson log-likelihood plus a Gaussian acceleration prior whose scale was estimated from second differences of training flights, warm-started from näıve Bayes with continuity and refined by L-BFGS then damped Newton steps over four restarts.

### 5.9 Encoding baselines

All GLMs used a raised-cosine place basis over *z*-scored position with a log link function, fit independently per session. Basis resolutions were anisotropic and selected by grid search on validation FDE— (6, 6, 3) for the place GLM and (4, 4, 2) for the trajectory-specific GLM — as coarse bases generalized markedly better than dense ones. The trajectory-specific GLM replaced the single readout with one readout per trajectory cluster plus a catch-all; clusters were derived from training flights only, retaining those with *≥* 3 flights, with held-out flights assigned to their nearest training flight. Lagged variants concatenate the same basis evaluated at positions from *−*1.0 to +1.0 s in 0.2 s steps. History, coupling and acausal variants add log-raised-cosine temporal filters [66]: six history bases over lags 0.1–1.0 s, four coupling bases over lags 0–1.0 s from all other neurons with the diagonal masked, and an acausal self filter with independent past and future weights. Coupling models carried a group lasso over off-diagonal filters whose strength was searched jointly with weight decay, and coupling parameters used a lower learning rate and no weight decay. The “MLP only” model is the twin with attention ablated to per-timestep feed-forward blocks. GLMs were fit by AdamW with early stopping on median validation FDE, with weight decay tuned by 10 Optuna trials.

### 5.10 Surrogate-trained controls

For the trial shuffle, flights within a cluster were warped to a common length and each neuron’s spike train permuted independently across flights of that cluster. For the synthetic surrogates, a smoothed 3-D rate map was fit per session — or per session and cluster — on training flights, and training spikes were replaced, per neuron independently, by draws from the empirical count distribution conditioned on spatial bin. Empirical rather than Poisson resampling preserves the real over-dispersion, while across-neuron simultaneity and within-flight temporal structure are destroyed. Every model was retrained on the surrogate with its architecture and hyperparameter protocol unchanged, including re-searching GLM weight decay and coupling penalty on the surrogate data. Validation and test sets remained real, non-surrogate data throughout (Fig. S2).

### 5.11 Context window and training-set size

Models with different context window sizes were trained with equal amounts of data by re-tiling every flight into non-overlapping windows of that length in both training and test loaders (Fig. S3). Each window received a 50-trial Optuna search of 300 epochs with *λ* pinned so all windows share one loss balance, followed by a 1,000-epoch refit. For decoding, a flight’s tiles were reordered and concatenated back into whole flights before the Fŕechet distance was computed. The ‘Full’ point in Fig. S3A was retrained under an identical protocol, and so is not the model used elsewhere in the paper. For the scaling analysis, twins were trained on 1, 2, 4 or 8 sessions under the production recipe, with 10 stratified subsets per size constructed so every session appears at least once; a session’s plotted value is the median over every subset containing it, and therefore averages across independently trained models.

### 5.12 Flight axis and motif segmentation

The principal flight axis of each room (Fig. S4) was measured on training flights only. Flights under 8 bins or 1 m of path were excluded, and each remaining flight was resampled to 50 points equally spaced in arc length. A flight’s own axis is the leading eigenvector of its covariance, and because that sign is arbitrary all derived quantities are sign-folded.

Each room’s mean axis is the leading eigenvector of the orientation tensor 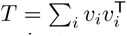 over its training flights, signed to +*x*. The orientation tensor is the standard sign-invariant aggregator of undirected axes, and is akin to the doubled-angle (semicircular) mean, but in three dimensions.

Axial progress *s*(*t*) is the projection of position onto that axis, centered on the environment’s mean flight centroid and smoothed by a centered 3-bin moving average. Each flight was labeled with at most one direction switch, which has a closed form: for a split after bin *k* the two legs advance by *a* and *b*, a switch is admissible only when *ab <* 0, and the gain over a single motif is 2 min(*|a|, |b|*), the overshoot of the turning point past the nearer endpoint. The optimal split was accepted only when that overshoot reached max(0.5 m, 0.25*×* the flight’s axial range), with each leg spanning at least 3 bins and 15% of the flight.

### 5.13 Latent extraction and representational spaces

Latent analyses were performed on a single evaluation pass over the test split of all 17 sessions. Two inference passes were executed per flight, with latent states extracted from the *neural* half of the token sequence to map both passes into a unified latent space: the encode pass masked all neural tokens while keeping behavioral tokens visible, whereas the decode pass left observed spike tokens visible while masking behavioral tokens. Encode and decode neural latents were concatenated before computing PCA. Latents are 64-dimensional and per-timestep, with no averaging or smoothing. Each flight was assigned a route label according to its starting and ending positions, where endpoints within 0.6 m of a feeder or perch were designated as reward or perch sites, respectively; flights failing this spatial threshold remained unclassified. Physical 3D spatial coordinates were additionally *z*-scored within each session, so room offset and scale do not dominate distances. For Hausdorff distance calculations and motif decoding, CEBRA and Twin latents were evaluated at their native dimensionality, using PCA strictly for visualization. We computed RSA at the motif level, splitting flights into segments if they contained two motifs. From panel C onward, we use only Reward*→* Reward, Reward*→*Perch, or Perch*→*Reward flights in Fig. 2.

### 5.14 RSA quantification

Each flight segment was represented as a full, variable-length point cloud without temporal normalization, resampling or time-averaging, allowing direct pairwise comparison between segments of distinct durations. For two point sets *A* and *B*, the directed Hausdorff distance was defined as

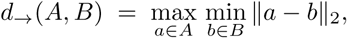

representing the maximum distance from any point in *A* to its nearest neighbor in *B*. Throughout this study, we evaluated the symmetric Hausdorff distance, defined as

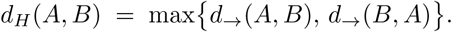

As a metric on compact sets, *d_H_* requires no point-to-point correspondence and is small if and only if every point in each set lies close to at least one point in the other. Consequently, two flight segments are judged as similar only if they traverse the same region across their entire trajectory rather than merely matching on average. Within each space, the resulting segment-by-segment matrix was *z*-scored using the mean and standard deviation of its off-diagonal entries, bringing all four spaces onto a common scale..

The read-out is a leave-one-out contrast on that matrix. For each segment *p* in turn, the self-comparison is removed and the remaining segments are split into those sharing its motif and those of the opposite motif; the statistic is

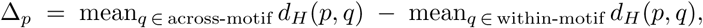

so Δ*_p_ >* 0 means segment *p* lies nearer other legs of its own motif than legs of the other motif. Every segment is held out exactly once and contributes one value. We admit only partners of a different route, so we measure generalization across routes. The same contrast was used for the dynamic-mode read-outs below.

Motif was also decoded from each space. Features are the time-averaged segment representations, taken from the same arrays the distance matrices were built from. The classifier is a two-step pipeline: a standard scaler fitted on the training fold only, followed by a support vector classifier with a radial basis function kernel, *C* = 1, and class weight = balanced to offset unequal motif counts. Again we cross-validated across routes by leave-one-route-out: each route is held out in turn and predicted by a classifier trained only on the remaining routes, a fold being used only when the training routes still carry both motifs. Train and test therefore never share a route, every segment is held out exactly once, and pooling the folds yields one out-of-sample prediction per segment. Physical XYZ is omitted from the decoding panels and is replaced by a raw-neural baseline on time-averaged spike counts; because the neuron space is session-specific, that baseline repeats the identical procedure within each session and pools the resulting predictions.

Performance is reported as balanced accuracy against a chance level of 0.5, with a class-stratified bootstrap 95% confidence interval over 1,000 resamples; the two plotted lines are the per-motif recalls, whose mean is that balanced accuracy. Spaces were compared by McNemar’s test over the segments both of them predicted, using the exact binomial form when fewer than 25 discordant pairs were available.

### 5.15 Neural latent decoding of kinematic variables

We trained linear regression models to predict kinematic variables — bat positions (X, Y, Z), velocity, acceleration, azimuth and elevation angles of heading, phase (relative time) — using the Twin encode latents as predictors. For comparison, we also used 3D physical positions and raw neural activity to train these models. Since the recorded unit count varied across sessions, cross-validation was conducted independently per session, with decoding performance evaluated using the R-squared (coefficient of determination) values. To account for their periodic nature, circular angular variables (azimuth and elevation) were converted to sine and cosine components prior to regression.

### 5.16 Dynamic mode decomposition

Dynamic mode decomposition [70–72] approximates a time series as a combination of dynamic modes, each consisting of a fixed spatial pattern evolving at a unique complex rate. We applied DMD to the Twin encode latent of individual qualified segments. Writing *L* for one segment’s latent, an *n × m* real matrix with *n* = 64 features and *m* time bins, the segment was first delay-embedded to depth *h* = 6 by stacking *h* copies of *L* shifted by one bin each,

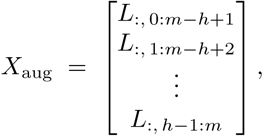

an *hn ×* (*m − h* + 1) matrix, which was split into paired, time-shifted snapshot matrices *X* = *X*_aug_[:, : *−*1] and *X^′^* = *X*_aug_[:, 1 :]. Stacking sacrifices *h −* 1 = 5 snapshots but lifts short segments out of the regime where the feature count exceeds the number of snapshots.

From the thin SVD *X* = *U* Σ*V ^∗^* at full rank, the projected linear dynamics matrix is given by *A* = *U ^∗^X^′^V* Σ*^−^*^1^. To preserve energy scaling across modes without relying on unit-normalized eigenvectors from standard eigensolvers, we performed a similarity transform *A*^^^ = Σ*^−^*^1^*^/^*^2^*A*Σ^1^*^/^*^2^. Eigendecomposition of *A*^^^*W*^^^ = *W*^^^ Λ yields eigenvalues Λ = diag(*λ_k_*) identical to the eigenvalues of *A*. We used the scaled eigenvectors *W* = Σ^1^*^/^*^2^*W*^^^ for further analysis. Exact modes were computed as Φ = *X^′^V* Σ*^−^*^1^*W* [71] with discrete eigenvalues *λ_k_*, whose continuous rates *ω_k_*= log *λ_k_/dt* define each mode’s frequency *|*Im *ω_k_|/*2*π* and its growth or decay Re *ω_k_*. Mode power was computed as 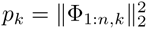, taken on the leading un-stacked block since the other *h −* 1 blocks are lagged copies of it.

Two features per segment were formed from this decomposition and converted to cosine distances, so that they enter the same across-minus-within-motif contrast as the Hausdorff matrices. The *temporal* feature is the power-weighted density of eigenvalues on the complex plane,

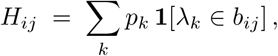

over a 19 *×* 19 grid of bins *b_ij_*spanning [*−*1.2, 1.2]^2^ in (Re *λ,* Im *λ*). The column of bins containing Im *λ* = 0 is set to zero, removing the non-oscillatory background so that the temporal feature is comparable with the spatial one, and *H* is then flattened and normalized to unit norm. Its dissimilarity is 1 *−* cos, the histograms being non-negative.

The *spatial* feature is a single mode’s pattern. Modes are ranked by descending *p_k_* after discarding those with *|*Im *λ_k_| ≤* 10*^−^*^9^ and, for each mode retained, consuming its complex-conjugate twin, so that the ranking enumerates distinct oscillations rather than raw eigenslots: a conjugate pair shares one real part and is not a second pattern, and the discarded real eigenvalues are the stationary background. The feature is Re Φ_1:*n,*_ *_k_∗* for the top-ranked *k*, normalized to unit length, and its dissimilarity is 1 *− |*cos*|* because a DMD mode is defined only up to a complex scalar, leaving the sign of its real part arbitrary.

### 5.17 Encode/decode and readout subspaces

Comparison dimensionality for the activation subspaces was set from the participation ratio, *d* = round(min(PR_enc_, PR_dec_)), each subspace being the top-*d* right singular vectors of its centered latent (Fig. S5). Readout matrices were row-stacked over sessions (neural) and environments (behavior) and compared to a dimension-permutation null, which randomizes only the correspondence between behavior-readout and latent dimensions.

Analyses compared subspaces of the 64-dimensional latent by the cosines of their principal angles. For orthonormal bases *A* and *B* these are the singular values of *A*^T^*B*, clipped to [0, 1]; a cosine of 1 means the two subspaces share a direction exactly, and 0 means that direction of one is orthogonal to all of the other. Components are reported in descending order, so the first is the best-aligned pair of directions and the last the worst. The null in Fig. S5B is 300 pairs of independent subspaces drawn uniformly at random, each obtained by QR factorization of a 64 *× d* Gaussian matrix, which fixes the chance level of alignment between two *d*-dimensional subspaces of a 64-dimensional space.

Fig. S5D asks how much of the behavior readout is reachable from the leading neural readout directions. Writing *V_k_* for the top-*k* right singular vectors of *W*_neu_, the energy of the behavior readout subspace inside that span is

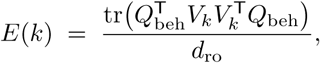

the mean squared cosine between the two subspaces, equivalently the fraction of the behavior readout subspace’s variance that survives projection onto *V_k_*. *E*(*k*) increases monotonically in *k* and is bounded by 1.

Panels Fig. S5C-D share one null: 300 relabelings of the latent dimensions, permuting the columns of *W*_beh_ by a single permutation applied to every row while the neural readout is held as observed.

A column permutation is orthogonal, so *W*_beh_*W* ^T^ — and with it every row norm, the cross-row structure and the entire singular spectrum — is left exactly intact, and only the correspondence between behavior-readout and latent dimensions is randomized.

### 5.18 Same-route, different-path flight pairs

Candidate pairs were within-session cluster pairs with at least five flights each whose mean trajectories both started and ended within 0.25 m of one another but whose discrete Fŕechet distance was at least 1.0 m. Four pairs survived across all 17 sessions, three of which are shown between Fig. 3 and Fig. S9. Paths 3 and 4 of Fig. 3 are the remaining clusters of the same route with at least three flights and at least one test flight, and enter display panels only. Normalized flight-progress maps used 40 equal bins of normalized cumulative path length, occupancy-normalized, with each spike assigned a progress value by interpolating its time onto the per-sample progress trace.

Place cells were selected on Path 1 alone. Its flights were shuffled once with a fixed seed and split 50/25/25, a split internal to this analysis and unrelated to the Twin’s; normalized flight-progress maps were then built separately on the first half and on the pooled held-out remainder. A cell had to satisfy three criteria: (i) the Pearson correlation between its training and held-out maps across the 40 progress bins exceeded the 97.5th percentile of 100 null values (one-sided *P <* 0.025), each null rebuilding the held-out map after circularly shifting every held-out flight’s spike times by an independent uniform offset within that flight’s own duration; (ii) the training map peaked at *≥* 1 Hz; and (iii) the correlation was finite, i.e. the cell was not silent on the held-out flights.

Goal heading is the allocentric azimuth minus the bearing to the reward feeder the flight lands on, left undefined within 0.10 m of the goal where that bearing is set by tracking noise alone. Heading-tuned cells were selected by the identical three-part criterion applied to each axis’s own 24-bin ring, with separate cell sets for azimuth and goal heading.

Each cell’s tuning was peak-normalized and centered across cells within each bin before computing population-vector correlations, and the bootstrap bands resample cells rather than diagonal elements, since adjacent bins are strongly autocorrelated. The near-versus-far null circularly rolls each cell’s Path-2 tuning independently. Peak-order tests use Spearman’s *ρ* on the normalized-flight-progress axis and the Fisher–Lee circular rank correlation on the heading axes, the latter being rank based and rotation invariant unlike a linear rank correlation on bin index.

### 5.19 Rate-map stability across paths

Rate maps were built separately for each geometric path of a route and compared pairwise, once in physical space and once in the first three principal components of the shared latent, fit per environment (Fig. 3K,L). A path needed at least 10 samples, and flights spanning less than 0.38 of their session’s bounding-box diagonal were excluded. Occupancy and spikes were binned, smoothed with a Gaussian with an SD of one bin and divided. The Wasserstein distance is EMD*/*diag, the exact earth-mover’s distance between the mass-normalized maps normalized by the longest transport the grid admits.

Because the transport cost is built from bin indices, an equal-count grid on an unequal box charges the same physical displacement differently along different axes. For this reason, grids were made cubic within each space and approximately matched on voxel count. A voxel was retained only where its smoothed occupancy reached 0.05 of the mass one sample deposits. The circular-shift null draws an independent within-flight offset per flight and neuron, so occupancy, path geometry and spike counts survive and only spike-to-place alignment is destroyed; a neuron’s null is the mean over 100 draws.

The bin-free estimator (Fig. 3M) compares the full latent against physical space without gridding either. Between two paths the Wasserstein distance is taken directly over the paths’ own sample points. Occupancy normalization is then automatic, nearest-neighbor estimation adapts to local density so no bandwidth in each space’s units need be chosen, and a sample-supported field cannot spill into unvisited regions, so no occupancy mask is required. Because raw values carry each space’s units and shrink with dimension, we do not compare this index across physical and latent spaces; the latent index was *z*-scored against its own circular-shift null. The reported latent is 15-dimensional, given that the latent’s participation ratio was 19.3 indoors and 14.8 outdoors.

### 5.20 Readout weights and subregion labels

Every neuron was summarized by its readout row normalized to unit norm, so only the direction of its readout was analyzed. The residual readout orthogonalizes each row against the population firing-rate axis *W* ^T^(*f −f*) and renormalizes. The firing-rate null in Fig. 4A projects the same readout matrix onto 100 random unit directions per session and takes the median absolute correlation with firing rate. Hierarchical clustering was cut at 12 clusters with only clusters of at least 200 members retained, five surviving; cutting fine and filtering by size was preferred to a small fixed *k*, because the residuals over-split into singletons while a small *k* merged distinct groups. No stability resampling was applied. Only reasonably well-fit neurons (Poisson FDE *>* 0) were used in Fig. 4, leaving 2,147 of 3,080 neurons.

CA1 and CA3 labels were assigned geometrically from unit depths, pooling by bat and probe: the depth histogram was smoothed (*σ* = 200 *µ*m) and up to two peaks detected with a minimum separation of 450 *µ*m, each defining a disjoint *±*300 *µ*m window, with the more superficial clump labeled CA1. Neurons outside both windows were not assigned a subregion, and subregion analyses required at least 10 labeled neurons per subregion per session. These assignments corresponded visually with the probe tracks from histology.

### 5.21 Tuning, attribution and similarity matrices

Rate maps for Fig. 4 were computed on the identical train split used for the Twin and GLM. We binned spikes in 0.3 m bins, excluding bins visited fewer than three times in either map, and computed each neuron’s cross-validated spatial information. This was defined as the log-likelihood of the test-split spikes under the train-split rate map minus their log-likelihood under a spatially uniform rate equal to the test set mean, normalized to bits per held out spike. Negative values indicate that the flat rate is a better predictor of held-out spiking. For attribution, the full Jacobian of predicted rate with respect to the position sequence was computed per test flight and attribution taken as its norm over the three position coordinates; flights were truncated to their first 200 valid bins, and the future and past sums accumulated across all test flights of a session before forming each neuron’s index. We also confirmed that attribution index differed across clusters even after controlling for differences in place field peak as a function of normalized flight progress (not shown).

Four neuron-by-neuron matrices were built per session in a common neuron order: cosine similarity of residual readouts; rate-map correlation; coupling from a separately fitted Poisson GLM with a place basis, history filter and coupling filter [66], taken as the time integral of the fitted filter and symmetrized; and the correlation of raw 0.1 s counts over all non-flight bins, that is, every bin in which the tracked flight indicator was zero. Four sessions with sufficient flights had a same-day sleep recording which furnished a fifth matrix; flight units were matched to sleep units with UnitMatch [108], run externally on the Kilosort4 output of both recordings from raw mean waveforms at a match probability threshold of 0.5, retaining the highest-probability partner per flight unit.

In the multiple regression on similarity matrices, non-finite pairs were removed by greedily dropping whole rows and columns until every surviving pair was finite in all four matrices, since the permutation null relabels neurons; sessions with fewer than 30 surviving pairs were dropped. The Freedman–Lane null fits the reduced model, reshapes its residuals into a symmetric matrix, relabels that matrix by a single permutation applied jointly to rows and columns, adds the permuted residuals back to the reduced fit and recomputes the statistic, so the exchangeable unit is the neuron and the pair-level dependence structure is preserved.

The subregion classifier used a standardized RBF-kernel support-vector machine with balanced class weights, the penalty tuned in an inner stratified cross-validation and balanced accuracy averaged over an outer repeated stratified loop of 20 repeats, so the reported value does not hinge on one split. For Fig. S10, the orthogonality chance level draws the same number of random unit vectors in 64 dimensions as the session has neurons.

## Acknowledgements

We thank members of the M.M.Y. (NeuroBat) laboratory for helpful discussions and the staff of the Office of Laboratory Animal Care for support with animal husbandry and care. This work was supported by a Jane Coffin Childs Fellowship (to A.S.L), and the Howard Hughes Medical Institute (to M.M.Y.). A.S.T acknowledges the support of NIH R01 NS113890 and the James Fickel Enigma Project Fund. The funders had no role in study design, data collection and analysis, decision to publish, or preparation of the manuscript.

## Author contributions

M.M.Y. and A.S.T. conceived and supervised the project. A.S.L. and Y.Q. developed the computational framework and performed the analyses, with contributions from Y.L. A.F. helped curate the dataset and advised on its interpretation. K.C.K. contributed to data interpretation. A.S.L. and Y.Q. prepared the figures and wrote the initial draft of the manuscript. M.M.Y. and A.S.T. revised the manuscript with input from all authors.

---

**Fig. S1.**
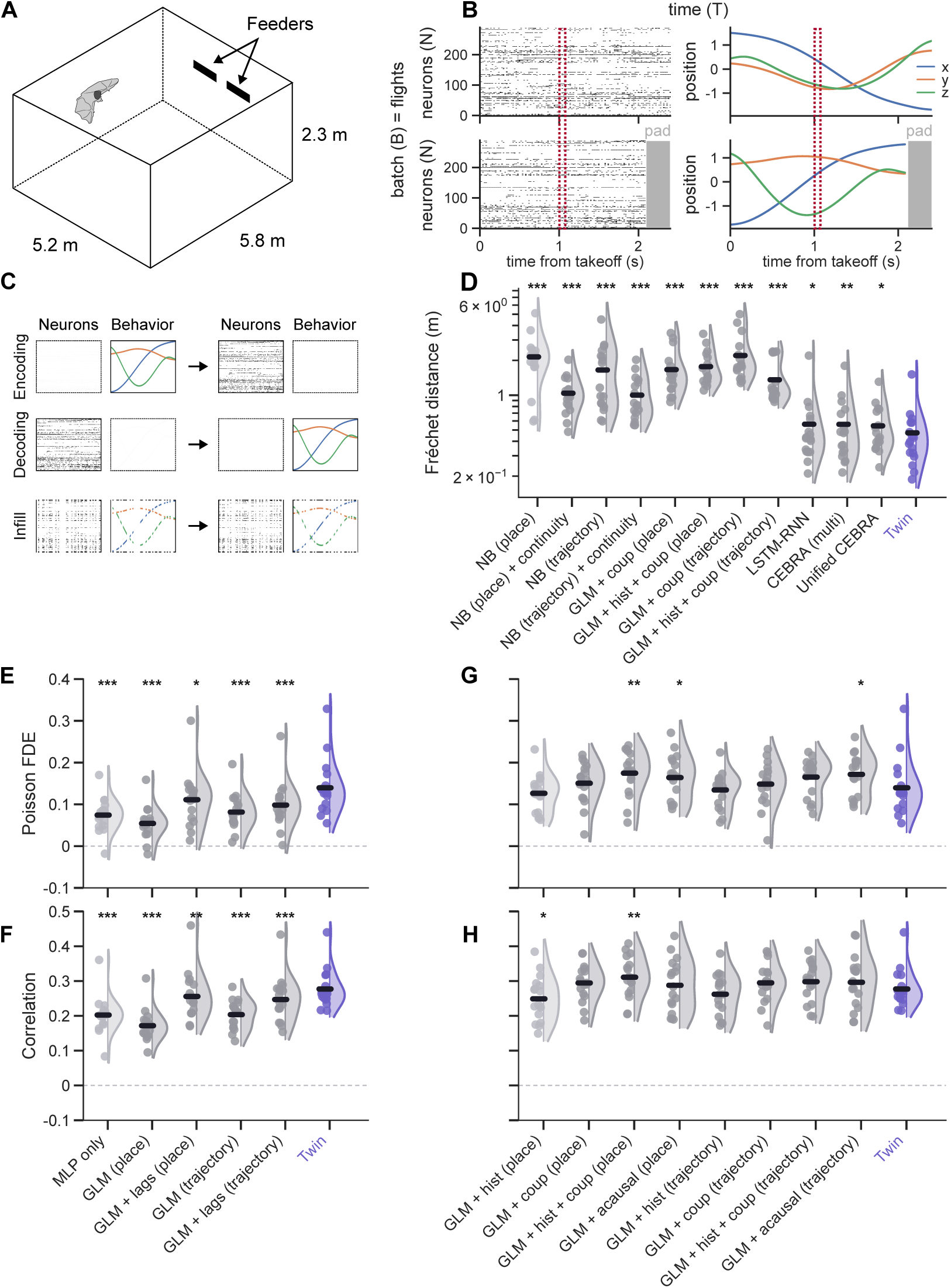
Model setup and full model ladders. (A) Schematic layout of indoor flight room with freely-moving recording. (B) Schematic input data for the Twin model. Single-unit spike counts and 3-D position were chunked into individual flights, binned at 100 ms, padded to the length of the longest flight, and input to the model with shape (B, T, N) and (B, T, 3), where the batch dimension *B* = number of flights, *T* = number of time bins, *N* = number of neurons, and 3 = number of spatial dimensions. (C). Each batch was assigned one of three masking strategies: “Encoding” (in which neural data was fully masked and used as the prediction target), “Decoding” (in which behavioral data was fully masked and used as the prediction target), and “Infill” (in which the same random time steps from both neural activity and behavior were masked and used as the prediction target). (D) Per-flight 3-D Fŕechet distance (lower is better) for the decoder ladder. GLMs were inverted by gradient descent over position to maximize Poisson likelihood + a smoothness prior, producing correlation-aware decoders (see Methods). Left to right, each versus the Twin (paired two-sided Wilcoxon signed-rank on session medians, *n* = 17 sessions): ‘NB (place)’ (*P* = 1.53*×* 10*^−^*^5^), ‘NB (place) + continuity’ (*P* = 1.53*×* 10*^−^*^5^), ‘NB (trajectory)’ (*P* = 1.53*×* 10*^−^*^5^), ‘NB (trajectory) + continuity’ (*P* = 1.53 *×* 10*^−^*^5^), ‘GLM + coup (place)’ (*P* = 1.53 *×* 10*^−^*^5^), ‘GLM + hist + coup (place)’ (*P* = 1.53 *×* 10*^−^*^5^), ‘GLM + coup (trajectory)’ (*P* = 1.53 *×* 10*^−^*^5^), ‘GLM + hist + coup (trajectory)’ (*P* = 1.53 *×* 10*^−^*^5^), ‘LSTM-RNN’ (*P* = 0.0129), ‘multi-session CEBRA’ (*P* = 0.00168), ‘Unified CEBRA’ (*P* = 0.0448), and our Twin. (E) Poisson FDE for models that use only behavior data for neural prediction. Each model versus the Twin (paired two-sided Wilcoxon signed-rank on session medians, *n* = 17 sessions): ‘MLP only’ (*P* = 7.63 *×* 10*^−^*^5^), ‘GLM (place)’ (*P* = 1.53 *×* 10*^−^*^5^), ‘GLM + lags (place)’ (*P* = 0.0110), ‘GLM (trajectory)’ (*P* = 3.05 *×* 10*^−^*^5^), ‘GLM + lags (trajectory)’ (*P* = 5.04 *×* 10*^−^*^4^). (F) Same as (E), but using Pearson correlation as metric: ‘MLP only’ (*P* = 1.53 *×* 10*^−^*^5^), ‘GLM (place)’ (*P* = 1.53 *×* 10*^−^*^5^), ‘GLM + lags (place)’ (*P* = 0.00168), ‘GLM (trajectory)’ (*P* = 1.53 *×* 10*^−^*^5^), ‘GLM + lags (trajectory)’ (*P* = 3.81 *×* 10*^−^*^4^). (G-H), same as (E-F), but for GLMs which additionally condition on spike history, cross-neuron coupling, and/or future spiking. (G) Poisson FDE: ‘GLM + hist (place)’ (*P* = 0.207), ‘GLM + coup (place)’ (*P* = 0.263), ‘GLM + hist + coup (place)’ (*P* = 0.00557), ‘GLM + acausal (place)’ (*P* = 0.0395), ‘GLM + hist (trajectory)’ (*P* = 1.00), ‘GLM + coup (trajectory)’ (*P* = 0.159), ‘GLM + hist + coup (trajectory)’ (*P* = 0.0714), ‘GLM + acausal (trajectory)’ (*P* = 0.0202). (H) Pearson correlation: ‘GLM + hist (place)’ (*P* = 0.0305), ‘GLM + coup (place)’ (*P* = 0.0984), ‘GLM + hist + coup (place)’ (*P* = 0.00557), ‘GLM + acausal (place)’ (*P* = 0.712), ‘GLM + hist (trajectory)’ (*P* = 0.159), ‘GLM + coup (trajectory)’ (*P* = 0.120), ‘GLM + hist + coup (trajectory)’ (*P* = 0.132), ‘GLM + acausal (trajectory)’ (*P* = 0.174). See also Fig. 1 and Figs. S2–S3.

**Fig. S2.**
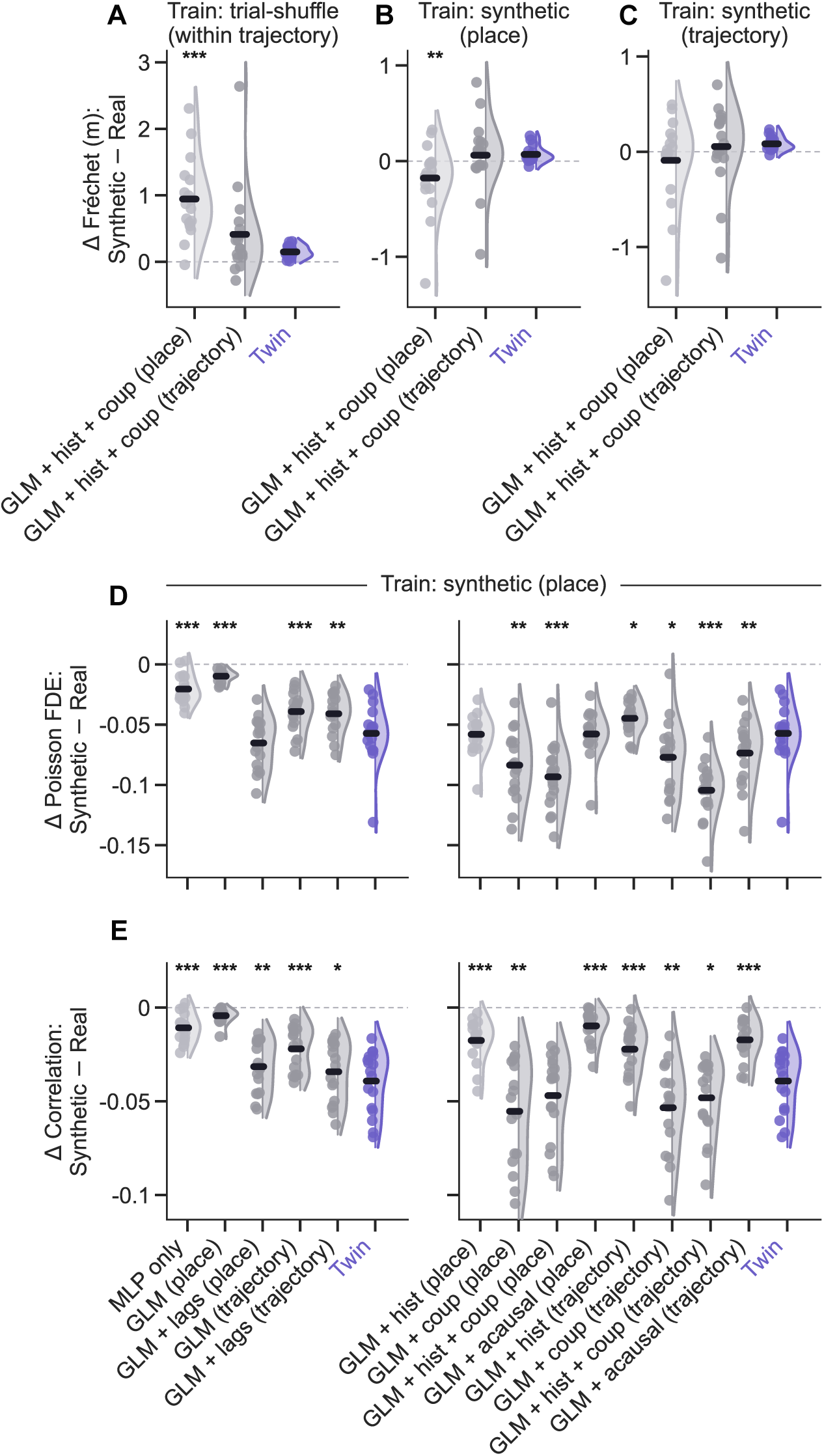
Surrogate-trained controls isolate the structure exploited by the digital Twin. All panels were trained on surrogate data as indicated and then evaluated on real, held-out test data. (A–C) Per-flight difference in 3-D Fŕechet distance (m) on surrogate versus real data. Three correlation-aware decoders of similar complexity and encoding performance (see Fig. S1) are compared: ‘GLM + hist + coup (place)’, ‘GLM + hist + coup (trajectory)’, and Twin, with the Wilcoxon test anchored on Twin. (A) Flights of the same trajectory cluster were randomly shuffled, breaking the “noise correlations” on which the ‘GLM + hist + coup (place)’ (*P* = 4.58 *×* 10*^−^*^5^) but not ‘(trajectory)’ (*P* = 0.145) model depend relative to the Twin. (B) Synthetic spikes were generated based on each recorded neuron’s binned and smoothed empirical rate map on training data, thereby eliminating not only noise correlations but also trajectory specificity. Now the ‘place GLM’, which matches the generative structure of the synthetic data, performs comparatively better than the Twin (*P* = 0.00665), while the ‘trajectory GLM’ does not (*P* = 0.712). (C) Same as (B), but synthetic spikes were generated in a trajectory-specific manner: ‘place GLM’ (*P* = 0.225) and ‘trajectory GLM’ (*P* = 0.678) vs. the Twin. (D) Per-neuron difference in Poisson fraction deviance explained between synthetic place-cell spikes, generated as in (B), and real data. The Twin is compared to the full model ladders from Fig. S1E,G: ‘MLP only’ (*P* = 3.81 *×* 10*^−^*^4^), ‘GLM (place)’ (*P* = 1.53 *×* 10*^−^*^5^), ‘GLM + lags (place)’ (*P* = 0.0569), ‘GLM (trajectory)’ (*P* = 8.39 *×* 10*^−^*^4^), ‘GLM + lags (trajectory)’ (*P* = 0.00464), ‘GLM + hist (place)’ (*P* = 0.890), ‘GLM + coup (place)’ (*P* = 0.00316), ‘GLM + hist + coup (place)’ (*P* = 3.05 *×* 10*^−^*^5^), ‘GLM + acausal (place)’ (*P* = 1.00), ‘GLM + hist (trajectory)’ (*P* = 0.0232), ‘GLM + coup (trajectory)’ (*P* = 0.0202), ‘GLM + hist + coup (trajectory)’ (*P* = 1.53 *×* 10*^−^*^5^), ‘GLM + acausal (trajectory)’ (*P* = 0.00934). The synthetic place cell spikes are least detrimental for models like ‘GLM (place)’ which are best matched to the data generation mechanism, and most detrimental for those that explicitly rely on cross-neuron coupling. (E) Same as (D) but for the difference in Pearson correlation: ‘MLP only’ (*P* = 1.07 *×* 10*^−^*^4^), ‘GLM (place)’ (*P* = 1.53 *×* 10*^−^*^5^), ‘GLM + lags (place)’ (*P* = 0.00790), ‘GLM (trajectory)’ (*P* = 7.63 *×* 10*^−^*^5^), ‘GLM + lags (trajectory)’ (*P* = 0.0110), ‘GLM + hist (place)’ (*P* = 1.07 *×* 10*^−^*^4^), ‘GLM + coup (place)’ (*P* = 0.00557), ‘GLM + hist + coup (place)’ (*P* = 0.120), ‘GLM + acausal (place)’ (*P* = 3.05 *×* 10*^−^*^5^), ‘GLM + hist (trajectory)’ (*P* = 1.07 *×* 10*^−^*^4^), ‘GLM + coup (trajectory)’ (*P* = 0.00316), ‘GLM + hist + coup (trajectory)’ (*P* = 0.0232), ‘GLM + acausal (trajectory)’ (*P* = 4.58 *×* 10*^−^*^5^). See also Fig. S1.

**Fig. S3.**
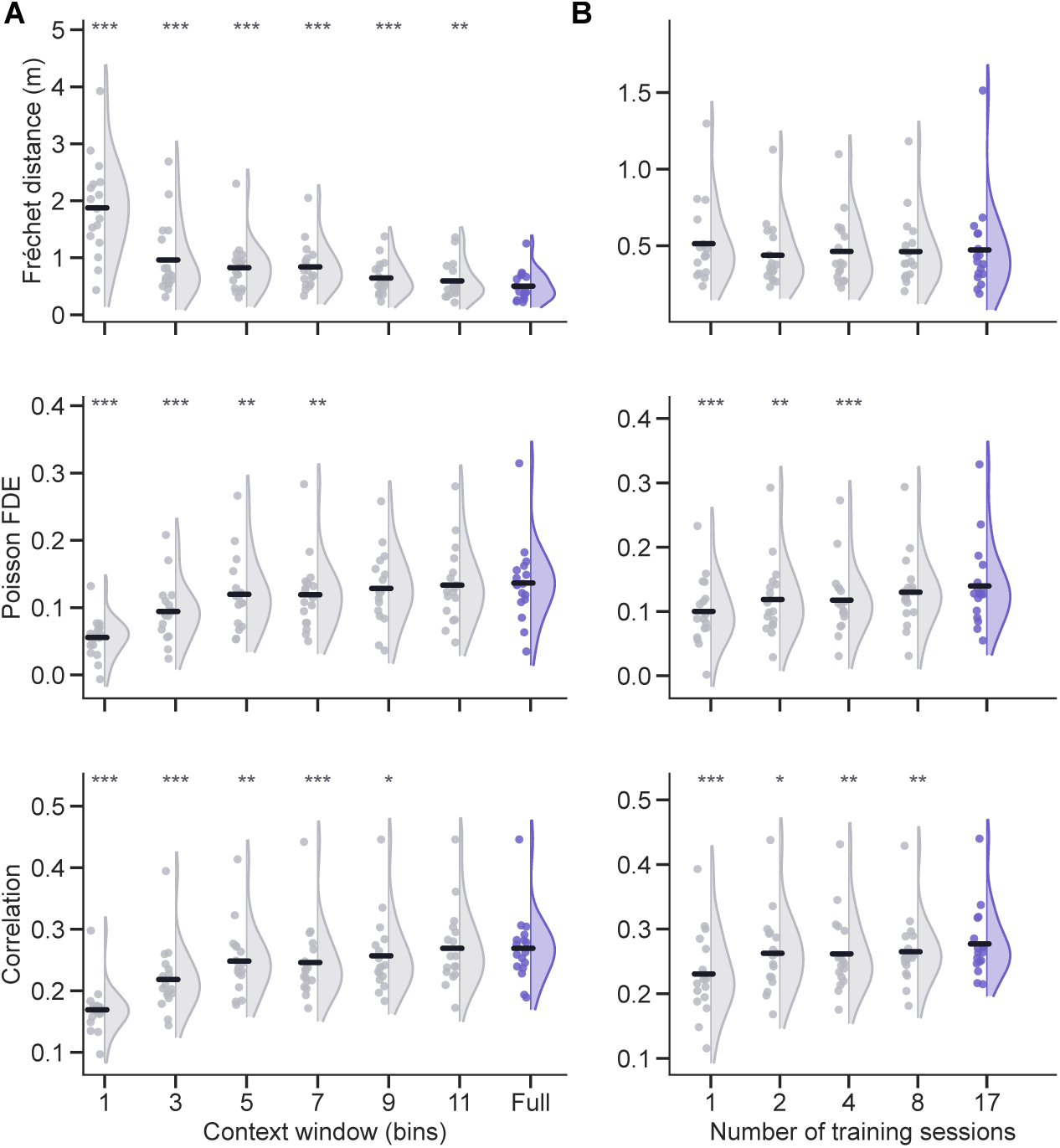
Scaling context window and training data improves digital Twin performance. (A) The Twin was retrained once per window size, holding the overall amount of data equivalent, and evaluated on 3-D Fŕechet distance between decoded and true flight paths (*top*), Poisson fraction deviance explained (*middle*), and Pearson correlation between predicted and observed spike counts (*bottom*). Every window size, ‘Full’ included, was hyperparameter-tuned and trained under one identical protocol, and decoded windows were re-assembled into whole flights so that all window sizes were scored on the same 278 test flights. Two-sided Wilcoxon signed-rank of each window vs. the full-context model gives *P* = 1.53 *×* 10*^−^*^5^ except at the longer windows: Fŕechet distance *P* = 2.58 *×* 10*^−^*^3^ (11 bins); Poisson FDE *P* = 6.65 *×* 10*^−^*^3^ (5 bins), *P* = 3.16 *×* 10*^−^*^3^ (7 bins), *P* = 0.174 (9 bins), *P* = 0.611 (11 bins); correlation *P* = 1.34 *×* 10*^−^*^3^ (5 bins), *P* = 3.82 *×* 10*^−^*^4^ (7 bins), *P* = 0.017 (9 bins), *P* = 0.611 (11 bins). Encoding is indistinguishable from full context by 9–11 bins, whereas the decoded path stays significantly worse at every window. (B) Same as (A), but as a function of the number of training sessions, where the 17-session model is the Twin used throughout. Encoding improves with more sessions: Poisson FDE *P* = 2.90 *×* 10*^−^*^4^ (1), *P* = 4.64 *×* 10*^−^*^3^ (2), *P* = 5.04 *×* 10*^−^*^4^ (4), *P* = 0.064 (8); correlation *P* = 1.53 *×* 10*^−^*^5^ (1), *P* = 0.015 (2), *P* = 2.09 *×* 10*^−^*^3^ (4), *P* = 1.07 *×* 10*^−^*^3^ (8) — whereas decoding Fŕechet distance shows only a trend at the smallest set (*P* = 0.071 at 1; *P* = 0.329, 0.678, 0.782 at 2, 4, 8; n.s.). All models are evaluated on the same per-session test split as the Full model. See also Fig. 1.

**Fig. S4.**
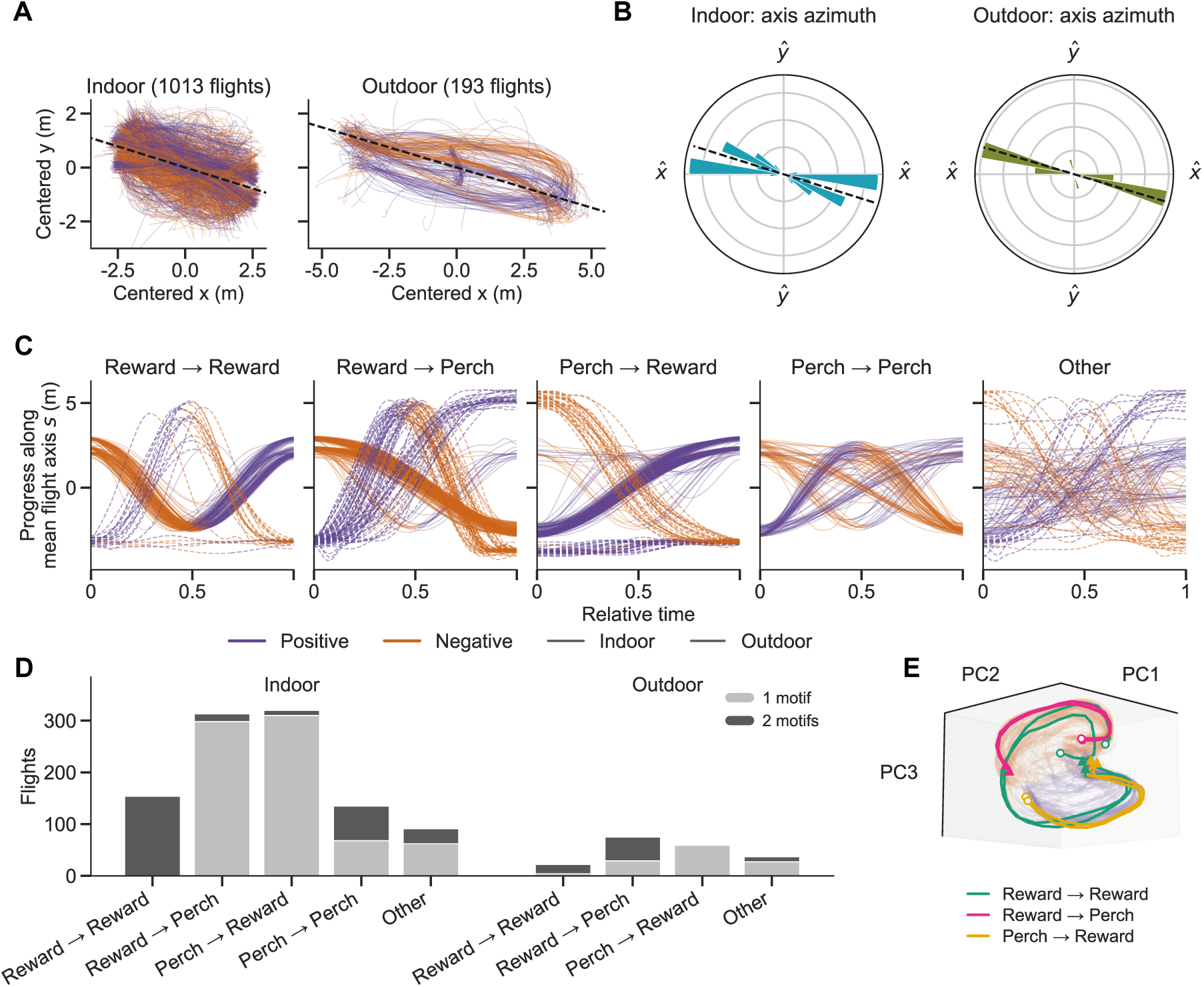
Flight in both rooms is organized along a principal axis, which defines motifs. (A) Every training flight seen from above, each translated onto its own centroid so that only its shape is shown, for the indoor room (*n* = 1,013 flights, 13 sessions, 4 bats) and the outdoor enclosure (*n* = 193 flights, 4 sessions, 2 bats). Each flight is cut into at most two motifs — travel along (+) or against (*−*) the room’s mean flight axis — and colored accordingly. Dashed line, the fitted axis of (B), drawn through the origin. (B) Circular histograms of orientation of the per-flight principal axis, obtained by PCA. Dashed line, the room’s fitted axis. (C) Signed progress *s*(*t*) along each room’s fitted axis for every flight in (A), one column per route and in normalized flight progress, colored by motif; indoor solid, outdoor dashed, columns sharing one y axis. The motif boundary is the turning point that maximizes the overshoot past the nearer endpoint of *s*, accepted only when that overshoot clears 0.5 m and 25% of the flight’s axial range. (D) Number of motifs per flight, by route and environment. (E) The same segmentation seen in the Twin’s encode latent, in the shared encode/decode PCA basis and at the viewing angle used for the Twin latent in Fig. 2G. Every held-out indoor flight is drawn faintly in its motif color, which splits the latent into two lobes; overlaid are the two flights per route (route palette) whose motifs sit deepest along the axis joining the two lobe centroids, one flight per session. Reward *→* perch and perch *→* reward flights are single-motif by construction and stay within one lobe each, on opposite sides; a reward *→* reward loop is one motif of each and has to cross between them. Open circle, flight start; triangle, flight end. See also Fig. 2.

**Fig. S5.**
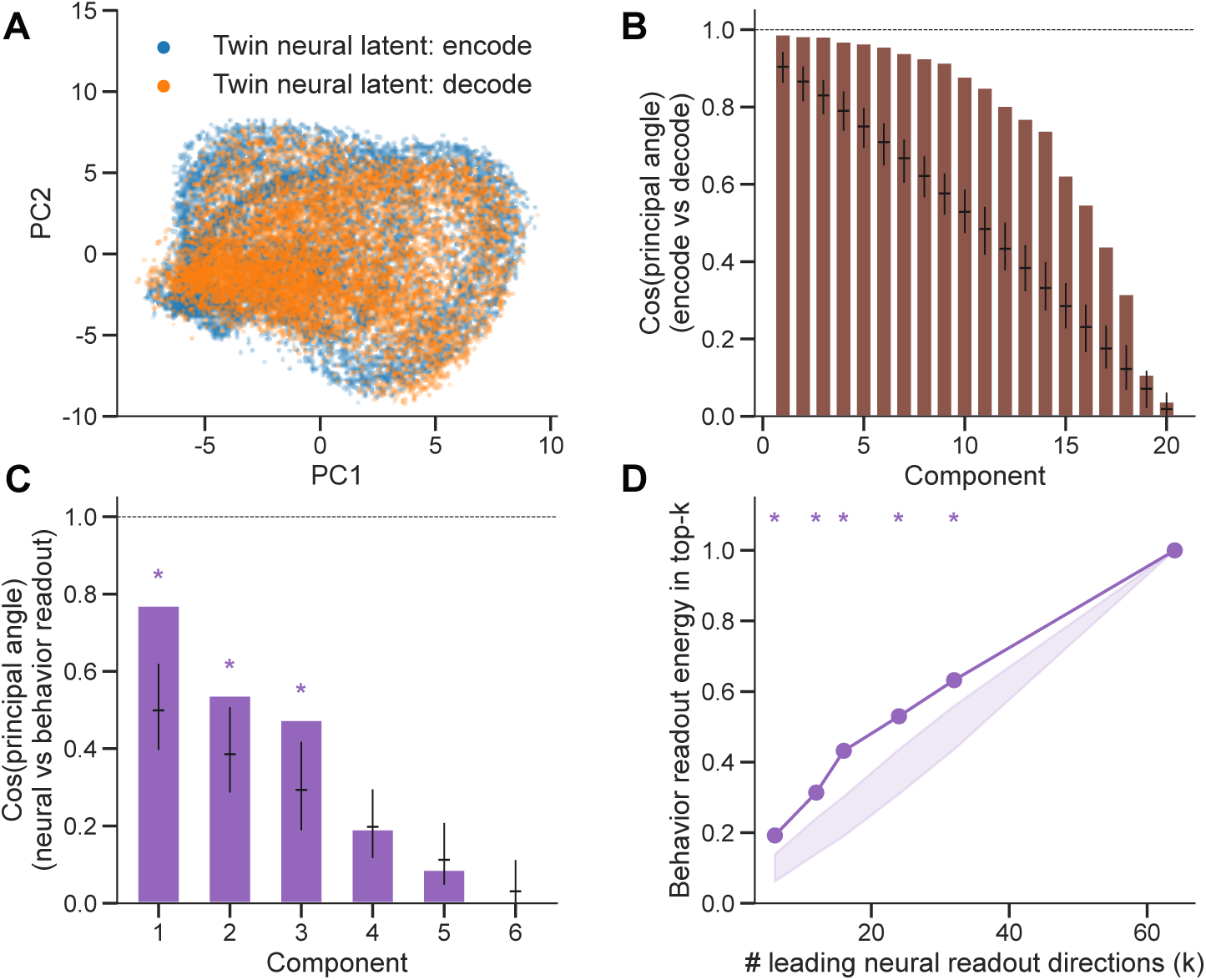
The digital Twin aligns encode and decode latents and readouts. (A) First two components of a single PCA fit to the row-concatenation of the encode and decode latents, so both share one basis. Latents are the pre-readout neural tokens on the encode and decode passes (*n* = 8,072 held-out timepoints per pass, pooled across 17 sessions). (B) Cosine of the top 20 principal angles between the encode and decode activation subspaces, 20 being the participation ratio of the latents (20.4 encode, 21.1 decode); bars = observed cosine, whiskers = 2.5–97.5% and tick = median of a null of principal angles between two independent random 20-dimensional subspaces of the 64-dimensional latent (*n* = 300). Dashed line at 1 = perfect alignment. (C) Cosine of the principal angles between the dominant neural and behavior readout directions, taking as many directions as the behavior readout rank (6); bars, whiskers, tick and dashed line as in (B), but against a dimension-permutation null: *n* = 300 relabelings of the 64 latent dimensions, permuting the columns of the behavior readout weight matrix by one permutation shared across rows while the neural readout is held as observed. Stars mark components falling outside the 2.5–97.5% interval of that null (two-sided permutation *P <* 0.05): components 1–3, *P* = 0.007, 0.033 and 0.007. (D) Fraction of the behavior readout subspace (its 6 leading right-singular directions) captured by the top-*k* neural readout directions, the latter taken over 3,080 neurons pooled across sessions; shaded band and stars as in (C), at *k* = 6, 12, 16, 24 and 32 (all *P* = 0.007). At *k* = 64 the fraction is 1 by construction. See also Fig. 2.

**Fig. S6.**
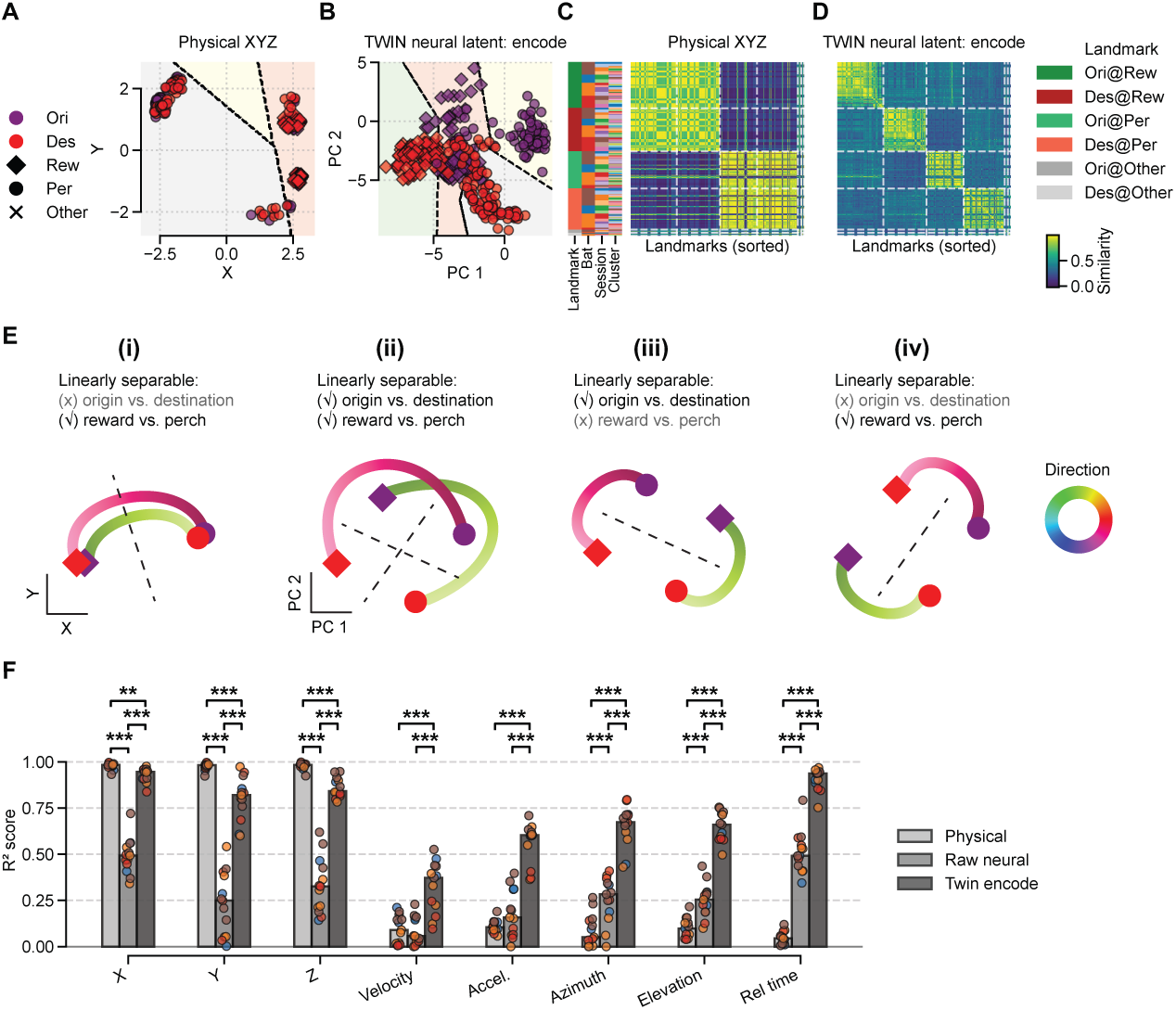
Representational similarity analysis (RSA) of landmark positions and neural latent decoding of kinematic variables. (A,B) 2D scatter plots of four landmark positions (flight origins and destinations at the reward and the perch) in physical space (A; X and Y as two axes) and population latent space (B; first two PCs) in the indoor environment. Purple and red indicate flight origins and destinations, respectively. Diamonds and circles indicate the reward and the perch, respectively. Dashed lines indicate decision boundaries from trained linear support vector machines (SVMs). We used Euclidean distance and cosine similarity as metrics to compute the similarity matrices in (A) and (B), respectively. We used the latent activations from Twin encode, which are followed by a linear transformation and a non-linear function to generate neural activity. (C,D) RSA matrices for landmarks (including flight origins or destinations at remaining positions) in 3D physical space (C) and 64D neural latent space (D). Matrix entries are hierarchically sorted by landmark identity, animal, experimental session, and flight cluster. White dashed lines indicate boundaries between landmark categories. Brighter hues on the color bar represent higher pairwise similarity scores. Note the four distinct block structures in latent space (D) compared to two blocks in physical space (C). (E) Schematics of different flight routes (reward-to-perch vs. perch-to-reward) in physical space (i) and latent space (ii), illustrating the motif structure in Fig. 2G and the landmark positions (A,B). Example trajectories for each route are colored distinctly, with color gradients representing heading azimuth angles. In physical space, reward and perch locations are linearly separable by design; however, flight origins and destinations overlap, and the two routes share similar geometry. Conversely, in the latent space identified by the Twin model, reward and perch locations remain linearly decodable, while flight origins and destinations become linearly separable. Furthermore, the two routes exhibit distinct geometrical structures. This arrangement is non-trivial, as demonstrated by two alternative hypotheses: (iii) where reward and perch locations are not linearly separable, and (iv) where origins and destinations are not linearly separable. Notably, no explicit optimization constraints were applied during model training; this manifold emerged naturally as a byproduct of linking bat flight positions with hippocampal activity. (F) Decoding accuracy (median R-squared score) for kinematic variables — bat positions (X, Y, Z), velocity, acceleration, azimuth and elevation angles of heading direction, phase (relative time) — using linear regression models trained on 3D physical positions (light gray), raw neural data (gray) and Twin encode latents (dark gray). Overlaid circle represents the score of individual session with different colors indicating distinct bats. Except the bat positions, the neural latent achieved superior decoding accuracy for all kinematic variables, exceeding 0.5 across seven variables (velocity proved challenging to decode across all feature spaces). Statistical comparisons: Two-sided Wilcoxon signed-rank tests across features: X position (*P* = 2.44 *×* 10*^−^*^4^ for physical vs. raw neural, *P* = 2.44 *×* 10*^−^*^4^ for raw neural vs. Twin encode, *P* = 1.22 *×* 10*^−^*^3^ for physical vs. Twin encode), Y position, elevation, and relative time (*P* = 2.44 *×* 10*^−^*^4^ for three pairwise comparisons), Z position (*P* = 2.44 *×* 10*^−^*^4^ for physical vs. raw neural, *P* = 2.44 *×* 10*^−^*^4^ for raw neural vs. Twin encode, *P* = 4.88 *×* 10*^−^*^4^ for physical vs. Twin encode), velocity (*P* = 5.42 *×* 10*^−^*^1^ for physical vs. raw neural, *P* = 2.44 *×* 10*^−^*^4^ for raw neural vs. Twin encode and for physical vs. Twin encode), acceleration (*P* = 1.10 *×* 10*^−^*^1^ for physical vs. raw neural, *P* = 2.44 *×* 10*^−^*^4^ for raw neural vs. Twin encode and for physical vs. Twin encode), azimuth(*P* = 4.88 *×* 10*^−^*^4^ for physical vs. raw neural, *P* = 2.44 *×* 10*^−^*^4^ for raw neural vs. Twin encode and for physical vs. Twin encode). See also Fig. 2.

**Fig. S7.**
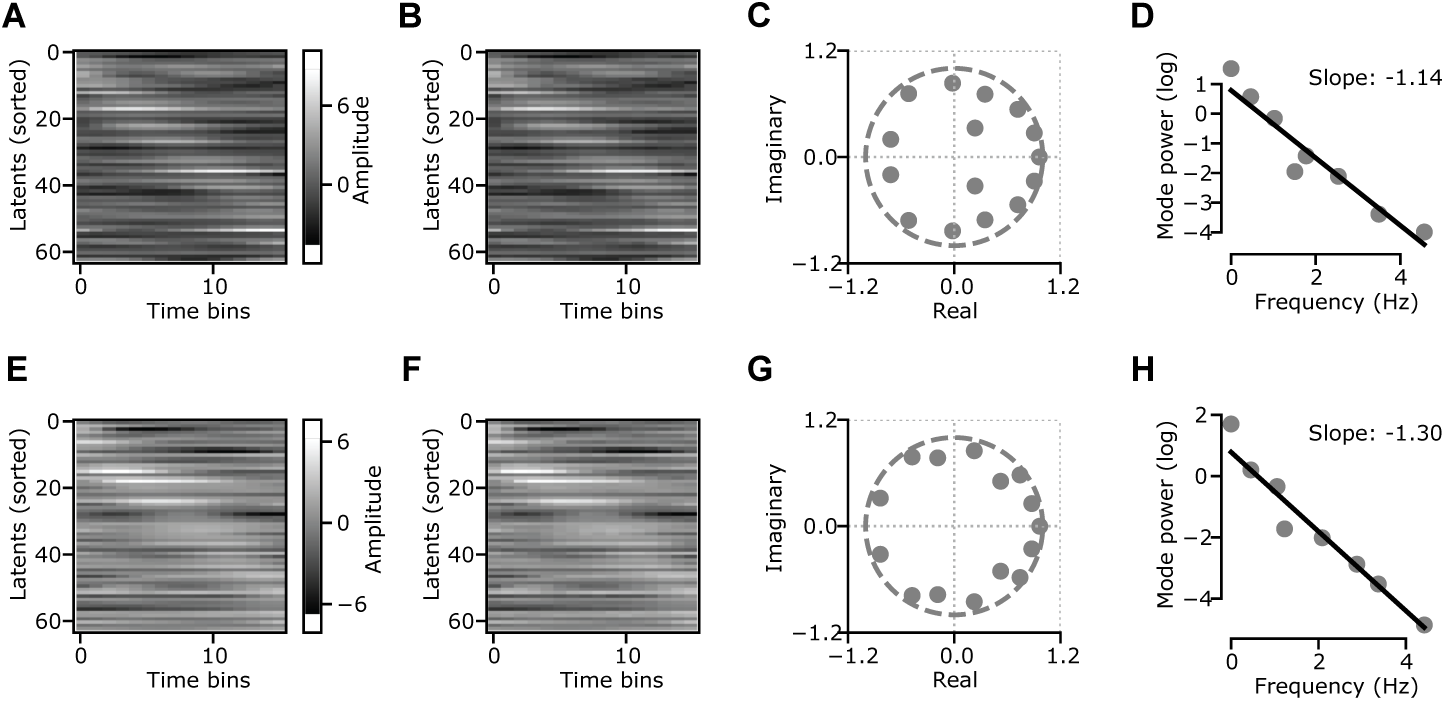
Dynamic mode decomposition (DMD) on exemplary flights. (A,B) Raw (A) and reconstructed (B) data for an example flight trajectory (origin at the reward and destination at the perch) in the Twin encode latent space. Latents are ordered by the time of peak activity across time bins on the raw data. (C) Eigenvalues (solid circles) on the complex plane for the example flight in A, shown relative to the unit circle (dashed line). (D) DMD power spectrum for the trajectory in A. Each circle corresponds to one DMD mode and the gray line represents a linear fit across modes. (E–H) Same as A–D, but for another exemplary flight with origin at the perch and destination at the reward. See also Fig. 2.

**Fig. S8.**
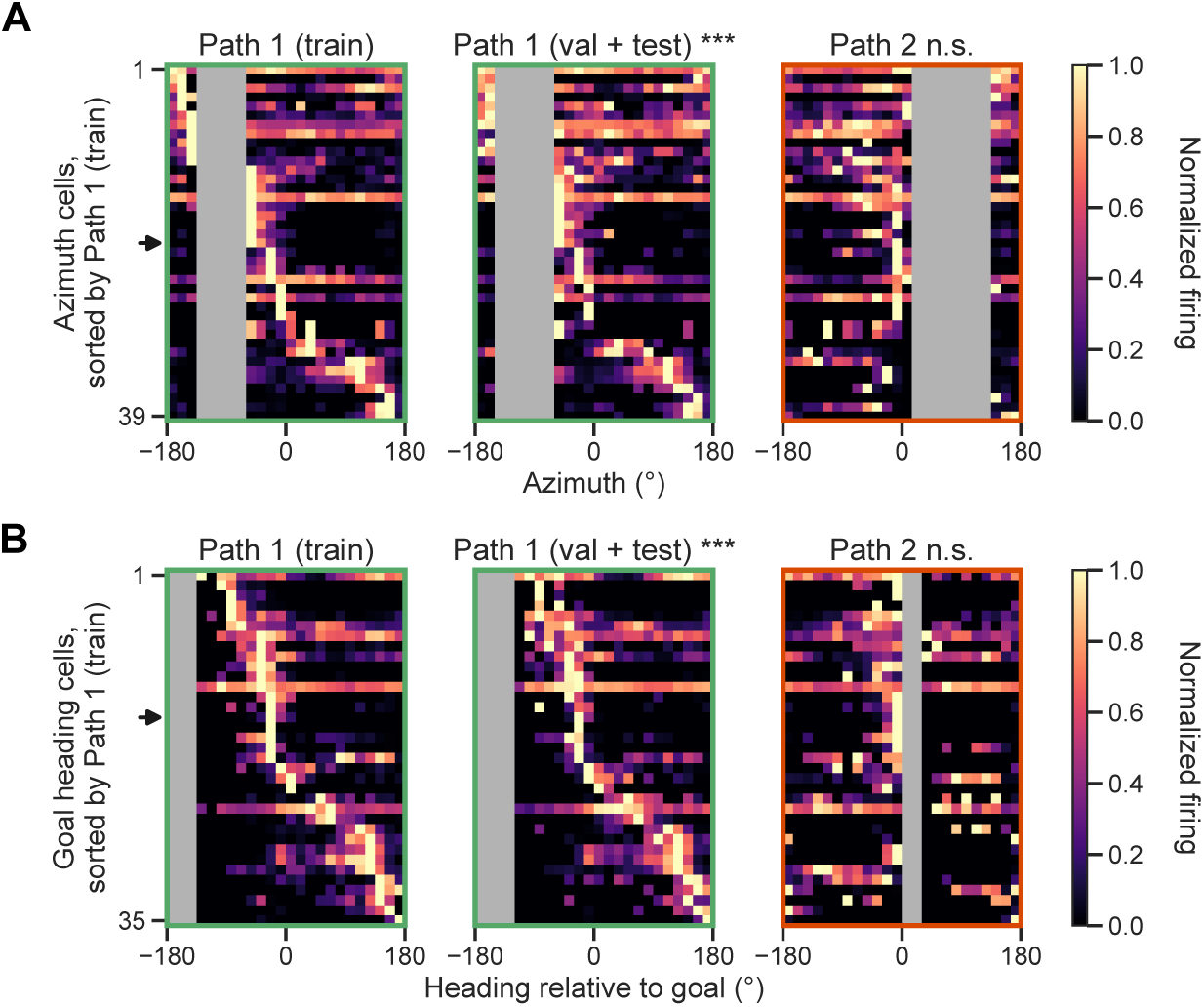
Generalization of place-cell sequences is not inherited from flight heading. (A) Firing-rate sequences on an azimuth axis (horizontal flight heading, 24 bins spanning *−*180 to 180*^◦^*) rather than normalized flight progress, on the Path 1 training flights (*left*), the held-out Path 1 validation + test flights (*middle*) and all Path 2 flights (*right*) of Fig. 3. Rows are the *n* = 39 azimuth-tuned cells. Grey bins were never occupied on those flights and are excluded from normalization and peak selection. The arrowhead marks the example unit of Fig. 3B,C. Panel titles carry the Fisher–Lee circular rank correlation between that panel’s azimuth-peak order and the Path-1-training order the rows are sorted by, against a label-permutation null. Azimuth-peak order is preserved within Path 1 (train vs. held-out *r* = 0.399, *P* = 5.00 *×* 10*^−^*^4^) but not between Path 1 and Path 2 (*r* = 0.037, *P* = 0.14) — the opposite of the normalized-flight-progress result in Fig. 3D, I. (B) As (A), for goal heading on *n* = 35 goal-heading-tuned cells. Both paths land on the same feeder, so this axis is common to them; nevertheless goal-heading-peak order is again preserved within Path 1 (*r* = 0.696, *P* = 5.00 *×* 10*^−^*^4^) and not between Path 1 and Path 2 (*r* = 0.049, *P* = 0.10). Permutation tests use 2,000 iterations. See also Fig. 3.

**Fig. S9.**
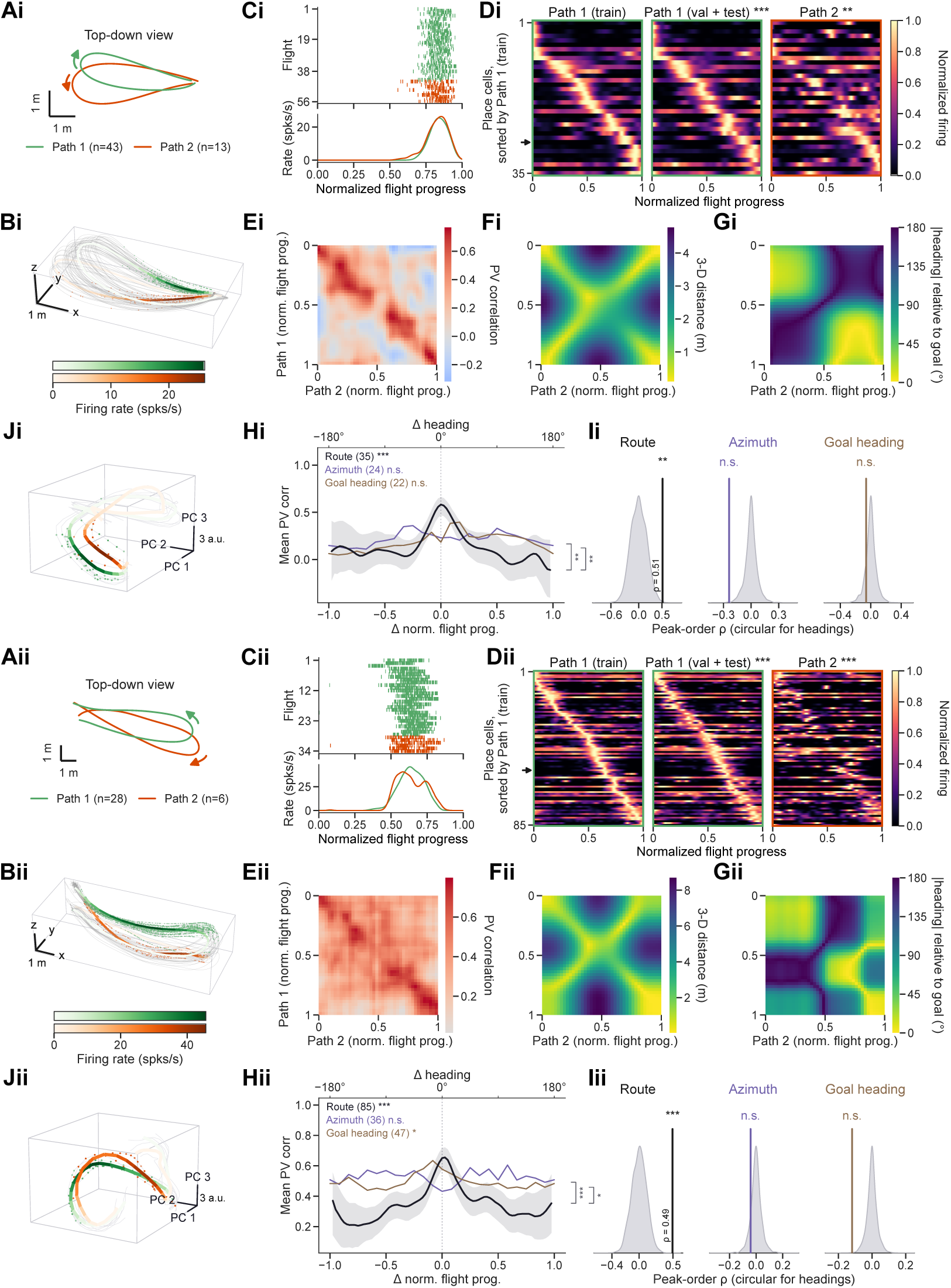
Additional examples of similar single-neuron tuning along geometrically different paths that share an origin and destination. Each subpanel as in Fig. 3A–J. Per example, the peak-order correlations of (D), the near-versus-far PV tests of (H) and the peak-order shuffle tests of (I): (i) bat 4, reward *→* reward, Path 1 = 43 flights vs. Path 2 = 13 flights, Fŕechet distance 1.97 m, *n* = 35 place cells (24 azimuth-, 22 goal-heading-tuned); peak-order *ρ* against the Path 1 training sort 0.765 held out (*P* = 5.00 *×* 10*^−^*^4^) and 0.580 on Path 2 (*P* = 1.50 *×* 10*^−^*^3^); PV near-vs-far +0.461 (*P* = 5.00 *×* 10*^−^*^4^) on normalized flight progress, +0.052 (*P* = 0.38) on azimuth and +0.143 (*P* = 0.071) on goal heading, the paired contrasts being +0.202 (*P* = 6.0 *×* 10*^−^*^3^) and +0.201 (*P* = 8.0 *×* 10*^−^*^3^); peak-order Spearman *ρ* = 0.513 (*P* = 2.00 *×* 10*^−^*^3^) between the held-out Path 1 flights and Path 2, against circular *ρ* = *−*0.238 (azimuth, *P* = 0.998) and *−*0.061 (goal heading, *P* = 0.85). (ii) bat 3, reward *→* perch, 28 vs. 6 flights, Fŕechet distance 2.36 m, *n* = 85 place cells (36 azimuth-, 47 goal-heading-tuned); peak-order *ρ* 0.747 held out and 0.429 on Path 2 (both *P* = 5.00 *×* 10*^−^*^4^); PV near-vs.far +0.323 (*P* = 5.00 *×* 10*^−^*^4^), *−*0.081 (*P* = 0.91) and +0.117 (*P* = 0.030), the paired contrasts being +0.209 (*P* = 1.00 *×* 10*^−^*^3^) and +0.099 (*P* = 0.032); peak-order Spearman *ρ* = 0.493 (*P* = 5.00 *×* 10*^−^*^4^), against circular *ρ* = *−*0.041 (*P* = 0.88) and *−*0.119 (*P* = 0.998). See also Fig. 3.

**Fig. S10.**
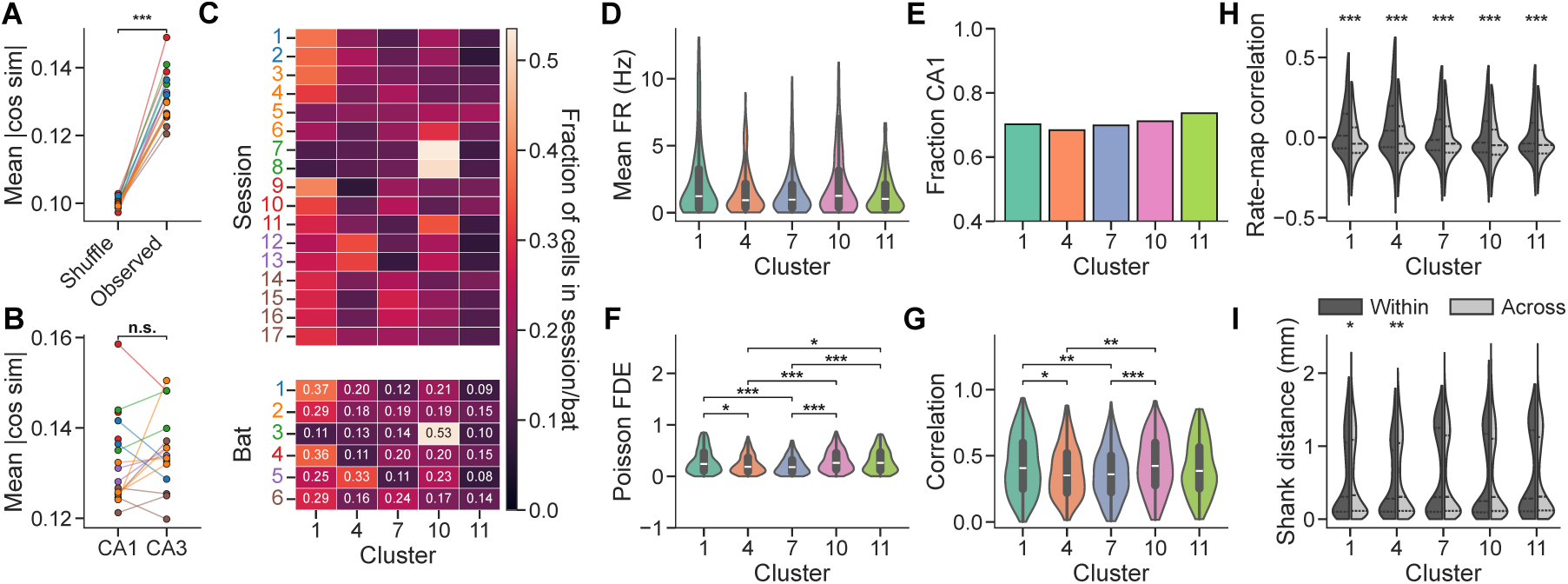
Residual readout cluster comparisons. (A) Residual readout weights are less orthogonal (higher within-session mean absolute cosine similarity) than random unit vectors of matched dimension. Each line is one session (colored by bat), plotted as the per-session mean; observed versus shuffle compared by two-sided paired Wilcoxon signed-rank test on per-session values (*P* = 1.53 *×* 10*^−^*^5^; *n* = 17 sessions). (B) Orthogonality (within-subregion mean absolute cosine similarity) does not differ between CA1 and CA3; two-sided paired Wilcoxon signed-rank, *P* = 0.330, n.s.; *n* = 15 sessions with *≥*10 neurons per subregion. (C) Fraction of each session’s (top) or each bat’s (bottom) well-fit neurons (Poisson FDE *>* 0), restricted to the five analyzed readout clusters, falling in each cluster. (D–G) Mean firing rate, fraction of CA1 neurons, Poisson FDE, and Pearson correlation per cluster. For (E), *χ*^2^ test of independence across clusters as omnibus, *P* = 0.775; two-sided Fisher’s exact test of each cluster’s CA1/CA3 split versus the pooled remaining clusters, all n.s. after Holm-Bonferroni correction across the five clusters, every adjusted *P* = 1.00). For other panels, Kruskal-Wallis omnibus across clusters; brackets mark every pairwise two-sided Mann-Whitney U comparison that is still significant after Holm-Bonferroni correction across the ten cluster pairs of that panel. Every P value quoted here for a comparison across clusters is Holm-adjusted within its own panel; omnibus tests are unadjusted. Mean FR: omnibus *P* = 0.155 (n.s.); no pair survives correction (smallest adjusted *P* = 0.127, 1 vs. 11). Poisson FDE: omnibus *P* = 2.91 *×* 10*^−^*^8^; significant pairs 1 vs. 4 *P* = 0.0156, 1 vs. 7 *P* = 1.53 *×* 10*^−^*^4^, 4 vs. 10 *P* = 2.55 *×* 10*^−^*^4^, 4 vs. 11 *P* = 0.0231, 7 vs. 10 *P* = 5.40*×*10*^−^*^7^, 7 vs. 11 *P* = 5.76*×*10*^−^*^4^. Pearson correlation: omnibus *P* = 3.87*×*10*^−^*^5^; significant pairs 1 vs. 4 *P* = 0.0320, 1 vs. 7 *P* = 0.00313, 4 vs. 10 *P* = 0.00387, 7 vs. 10 *P* = 2.60 *×* 10*^−^*^4^. (H–I) Rate-map correlation (H) and physical distance along the shank (I, mm) for neuron pairs within versus across cluster, shown per cluster, with a two-sided Mann–Whitney U comparing within versus across, Holm–Bonferroni corrected across the five clusters. (H) Cluster 1, *P* = 1.09 *×* 10*^−^*^184^; 4, *P* = 1.25 *×* 10*^−^*^132^; 7, *P* = 7.29 *×* 10*^−^*^26^; 10, *P* = 1.29 *×* 10*^−^*^51^; 11, *P* = 4.40 *×* 10*^−^*^17^. (I) Cluster 1, *P* = 0.0472; 4, *P* = 0.00784; 7, *P* = 0.767 (n.s.); 10, *P* = 0.0793 (n.s.); 11, *P* = 0.834 (n.s.).

**Table S1.** Auxiliary masking task sweep. The digital Twin model was trained from scratch (1,000 epochs, early stopping, 4 transformer layers, latent dimension *d* = 64, feed-forward width 64, learning rate 4.5*×*10*^−^*^3^, weight decay 3*×*10*^−^*^4^, decoding weight *λ* = 0.61, positive weight 0.84) on the encode and decode tasks plus every subset (0–4) of the four auxiliary self-supervised masking tasks within, cross (also called infill in the main text), forecast and forecast stim — 16 combinations in total. Scheme sampling weights are fixed at *w*_encode_ = 1.93 and *w*_decode_ = 2.83, and the total auxiliary-task probability is held constant across combinations by setting *w*_aux_ = 0.67*×*3*/n*_aux_. Rows are ranked by the validation joint score 0.5 (FDE*/*0.1) *−* 0.5 (err*/*0.3), which is the model-selection criterion; the decode error, encode FDE and joint score in the remaining columns are held-out test values, shown for reference only. The combination with the best validation score is shown in bold, and was used throughout the main text. For reference, the further fine-tuning the parameters of the cross model reaches a test joint score of 0.127.

| Rank | Auxiliary tasks | $n_{\text{aux}}$ | $w_{\text{aux}}$ | Decode err. (m) | Encode FDE | Joint (valid) | Joint (test) |
| --- | --- | --- | --- | --- | --- | --- | --- |
| 1 | **cross** | **1** | **2.010** | **0.2812** | **0.1093** | **0.0747** | **0.0781** |
| 2 | **within + cross + forecast + forecast_stim** | 4 | 0.503 | 0.2859 | 0.1139 | 0.0387 | 0.0929 |
| 3 | **within + forecast_stim** | 2 | 1.005 | 0.2878 | 0.1159 | 0.0309 | 0.0997 |
| 4 | **within + cross** | 2 | 1.005 | 0.2806 | 0.1099 | 0.0236 | 0.0815 |
| 5 | **cross + forecast_stim** | 2 | 1.005 | 0.2940 | 0.1084 | 0.0223 | 0.0518 |
| 6 | **cross + forecast** | 2 | 1.005 | 0.2900 | 0.1088 | 0.0108 | 0.0605 |
| 7 | **within + cross + forecast** | 3 | 0.670 | 0.2769 | 0.1129 | 0.0071 | 0.1032 |
| 8 | (none) | 0 | 0.000 | 0.2951 | 0.1029 | −0.0087 | 0.0228 |
| 9 | **within + forecast** | 2 | 1.005 | 0.3045 | 0.1052 | −0.0094 | 0.0185 |
| 10 | **within + cross + forecast_stim** | 3 | 0.670 | 0.2863 | 0.1152 | −0.0116 | 0.0989 |
| 11 | **within** | 1 | 2.010 | 0.3073 | 0.1071 | −0.0122 | 0.0231 |
| 12 | **within + forecast + forecast_stim** | 3 | 0.670 | 0.3016 | 0.1073 | −0.0466 | 0.0338 |
| 13 | **forecast_stim** | 1 | 2.010 | 0.3292 | 0.0967 | −0.0793 | −0.0651 |
| 14 | **forecast** | 1 | 2.010 | 0.3145 | 0.0975 | −0.0870 | −0.0368 |
| 15 | **forecast + forecast_stim** | 2 | 1.005 | 0.3057 | 0.0994 | −0.0871 | −0.0127 |
| 16 | **cross + forecast + forecast_stim** | 3 | 0.670 | 0.3585 | 0.1037 | −0.1199 | −0.0791 |

## References

1. Yartsev MM. Natural behavior is what you do, not just where you are. Nature Neuroscience. 2026;29(4):763–6.

2. Stevenson IH, Kording KP. How advances in neural recording affect data analysis. Nature Neuroscience. 2011;14(2):139–42.

3. Chen TW, Wardill TJ, Sun Y, Pulver SR, Renninger SL, Baohan A, et al. Ultrasensitive fluorescent proteins for imaging neuronal activity. Nature. 2013;499(7458):295–300.

4. Steinmetz NA, Aydin C, Lebedeva A, Okun M, Pachitariu M, Bauza M, et al. Neuropixels 2.0: A miniaturized high-density probe for stable, long-term brain recordings. Science. 2021;372(6539):eabf4588.

5. Anderson DJ, Perona P. Toward a science of computational ethology. Neuron. 2014;84(1):18–31.

6. Mathis A, Mamidanna P, Cury KM, Abe T, Murthy VN, Mathis MW, et al. DeepLabCut: markerless pose estimation of user-defined body parts with deep learning. Nature Neuroscience. 2018 Sep;21(9):1281–9.

7. Couzin ID, Heins C. Emerging technologies for behavioral research in changing environments. Trends in Ecology & Evolution. 2023;38(4):346–54.

8. Kong S, Deng F, Dong X, Liu C, Chen W, Wang Y, et al. Machine learning methods for studying latent neural activity dynamics. arXiv [csLG]. 2026 Jun.

9. Zhang C, Le Moing G, Koppula S, Rocco I, Momeni L, Xie J, et al. Efficiently reconstructing dynamic scenes one d4rt at a time. In: Proceedings of the IEEE/CVF Conference on Computer Vision and Pattern Recognition; 2026. p. 7382-92.

10. OpenAI, Achiam J, Adler S, Agarwal S, Ahmad L, Akkaya I, et al. GPT-4 Technical Report. arXiv [csCL]. 2023 Mar.

11. Jumper J, Evans R, Pritzel A, Green T, Figurnov M, Ronneberger O, et al. Highly accurate protein structure prediction with AlphaFold. Nature. 2021;596(7873):583–9.

12. Yamins DL, DiCarlo JJ. Using goal-driven deep learning models to understand sensory cortex. Nature Neuroscience. 2016;19(3):356–65.

13. Walker EY, Sinz FH, Cobos E, Muhammad T, Froudarakis E, Fahey PG, et al. Inception loops discover what excites neurons most using deep predictive models. Nature Neuroscience. 2019;22(12):2060–5.

14. Mathis MW, Mathis A. Joint modelling of brain and behaviour dynamics with artificial intelligence. Nature Reviews Neuroscience. 2026;27(2):87–100.

15. Ye J, Collinger J, Wehbe L, Gaunt R. Neural Data Transformer 2: Multi-context Pretraining for Neural Spiking Activity. In: Oh A, Naumann T, Globerson A, Saenko K, Hardt M, Levine S, editors. Advances in Neural Information Processing Systems. vol. 36. Curran Associates, Inc.; 2023. p. 80352–74.

16. Antoniades A, Yu Y, Canzano J, Wang W, Smith SL. Neuroformer: Multimodal and multitask generative pretraining for brain data. arXiv [q-bioNC]. 2023 Oct.

17. Azabou M, Arora V, Ganesh V, Mao X, Nachimuthu SB, Mendelson MJ, et al. A Unified, Scalable Framework for Neural Population Decoding. In: Thirty-seventh Conference on Neural Information Processing Systems; 2023. .

18. Azabou M, Pan KX, Arora V, Knight IJ, Dyer EL, Richards BA. Multi-session, multi-task neural decoding from distinct cell-types and brain regions. In: The Thirteenth International Conference on Learning Representations; 2025. .

19. Zhang Y, Wang Y, Azabou M, Andre A, Wang Z, Lyu H, et al. Neural Encoding and decoding at scale. arXiv [q-bioNC]. 2025 Apr.

20. Willeke KF, Turishcheva P, Gilbert A, Chakrabarty G, Bedel HA, Fahey PG, et al. OmniMouse: Scaling properties of multi-modal, multi-task Brain Models on 150B Neural Tokens. In: The Fourteenth International Conference on Learning Representations; 2026. .

21. Wang EY, Fahey PG, Ding Z, Papadopoulos S, Ponder K, Weis MA, et al. Foundation model of neural activity predicts response to new stimulus types. Nature. 2025 Apr;640(8058):470–7.

22. Ponce CR, Xiao W, Schade PF, Hartmann TS, Kreiman G, Livingstone MS. Evolving images for visual neurons using a deep generative network reveals coding principles and neuronal preferences. Cell. 2019;177(4):999–1009.

23. Bashivan P, Kar K, DiCarlo JJ. Neural population control via deep image synthesis. Science. 2019;364(6439):eaav9436.

24. Ding Z, Tran D, Ponder K, Ding Z, Froebe R, Ntanavara L, et al. Functional bipartite invariance in mouse primary visual cortex receptive fields. Nature Neuroscience. 2026;29(4):851–63.

25. Franke K, Willeke KF, Ponder K, Galdamez M, Zhou N, Muhammad T, et al. State-dependent pupil dilation rapidly shifts visual feature selectivity. Nature. 2022;610(7930):128–34.

26. Ding Z, Fahey PG, Papadopoulos S, Wang EY, Celii B, Papadopoulos C, et al. Functional connectomics reveals general wiring rule in mouse visual cortex. Nature. 2025;640(8058):459–69.

27. Fu J, Shrinivasan S, Baroni L, Ding Z, Fahey PG, Pierzchlewicz PA, et al. Statistics of natural scenes shape contextual modulation in the visual cortex. Neuron. 2026;114(13):2473–86.

28. Moser EI, Moser MB, McNaughton BL. Spatial representation in the hippocampal formation: a history. Nature Neuroscience. 2017 Oct;20(11):1448–64.

29. Eichenbaum H, Cohen NJ. Can we reconcile the declarative memory and spatial navigation views on hippocampal function? Neuron. 2014 Aug;83(4):764–70.

30. Lisman J, Buzaki G, Eichenbaum H, Nadel L, Ranganath C, Redish AD. Viewpoints: how the hippocampus contributes to memory, navigation and cognition. Nature Neuroscience. 2017 Oct;20(11):1434–47.

31. Whittington JCR, McCaffary D, Bakermans JJW, Behrens TEJ. How to build a cognitive map. Nature Neuroscience. 2022 Oct;25(10):1257–72.

32. O’Keefe J, Dostrovsky J. The hippocampus as a spatial map: preliminary evidence from unit activity in the freely-moving rat. Brain Research. 1971.

33. O’Keefe J, Nadel L. The hippocampus as a cognitive map. Oxford University Press; 1978.

34. McNaughton BL, Barnes CA, O’Keefe J. The contributions of position, direction, and velocity to single unit activity in the hippocampus of freely-moving rats. Exp Brain Res. 1983;52(1):41–9.

35. Muller RU, Bostock E, Taube JS, Kubie JL. On the directional firing properties of hippocampal place cells. Journal of Neuroscience. 1994 Dec;14(12):7235–51.

36. Markus EJ, Qin YL, Leonard B, Skaggs WE, McNaughton BL, Barnes CA. Interactions between location and task affect the spatial and directional firing of hippocampal neurons. Journal of Neuroscience. 1995;15(11):7079–94.

37. Wood ER, Dudchenko PA, Robitsek RJ, Eichenbaum H. Hippocampal neurons encode information about different types of memory episodes occurring in the same location. Neuron. 2000 Sep;27(3):623–33.

38. Frank LM, Brown EN, Wilson M. Trajectory encoding in the hippocampus and entorhinal cortex. Neuron. 2000 Jul;27(1):169–78.

39. Ferbinteanu J, Shapiro ML. Prospective and retrospective memory coding in the hippocampus. Neuron. 2003 Dec;40(6):1227–39.

40. Grieves RM, Wood ER, Dudchenko PA. Place cells on a maze encode routes rather than destinations. eLife. 2016 Jun;5:e15986.

41. Sarel A, Finkelstein A, Las L, Ulanovsky N. Vectorial representation of spatial goals in the hippocampus of bats. Science. 2017 Jan;355(6321):176–80.

42. Liberti III WA, Schmid TA, Forli A, Snyder M, Yartsev MM. A stable hippocampal code in freely flying bats. Nature. 2022;604(7904):98–103.

43. Ormond J, O’Keefe J. Hippocampal place cells have goal-oriented vector fields during navigation. Nature. 2022 Jul;607(7920):741–6.

44. Duvelle E, Grieves RM, van der Meer MAA. Temporal context and latent state inference in the hippocampal splitter signal. eLife. 2023 Jan;12(e82357):e82357.

45. Forli A, Fan W, Qi KK, Yartsev MM. Replay and representation dynamics in the hippocampus of freely flying bats. Nature. 2025;645(8082):974–80.

46. Zutshi I, Apostolelli A, Yang W, Zheng ZS, Dohi T, Balzani E, et al. Hippocampal neuronal activity is aligned with action plans. Nature. 2025 Mar;639(8053):153–61.

47. O’Keefe J. How the hippocampal cognitive map supports flexible navigation. Annu Rev Neurosci. 2025 Jul;48(1):331–44.

48. Maimon SR, Eliav T, Aljadeff J, Shalev A, Gronich Y, Finger NM, et al. Sparse-to-dense coding transformation between hippocampal areas CA3 and CA1. Nature. 2026 Jul;655(8125):1242–51.

49. Whittington JCR, Muller TH, Mark S, Barry C, Behrens TEJ. Generalisation of structural knowledge in the hippocampal-entorhinal system. arXiv [csAI]. 2018 May.

50. Whittington JCR, Muller TH, Mark S, Chen G, Barry C, Burgess N, et al. The Tolman-Eichenbaum Machine: Unifying Space and Relational Memory through Generalization in the Hippocampal Formation. Cell. 2020 Nov.

51. MacDonald CJ, Lepage KQ, Eden UT, Eichenbaum H. Hippocampal “time cells” bridge the gap in memory for discontiguous events. Neuron. 2011 Aug;71(4):737–49.

52. McKenzie S, Frank AJ, Kinsky NR, Porter B, Rivìere PD, Eichenbaum H. Hippocampal representation of related and opposing memories develop within distinct, hierarchically organized neural schemas. Neuron. 2014 Jul;83(1):202–15.

53. Constantinescu AO, O’Reilly JX, Behrens TEJ. Organizing conceptual knowledge in humans with a gridlike code. Science. 2016 Jun;352(6292):1464–8.

54. Aronov D, Nevers R, Tank DW. Mapping of a non-spatial dimension by the hippocampal-entorhinal circuit. Nature. 2017 Mar;543(7647):719–22.

55. Gauthier JL, Tank DW. A dedicated population for reward coding in the hippocampus. Neuron. 2018;99(1):179–93.

56. Schuck NW, Niv Y. Sequential replay of nonspatial task states in the human hippocampus. Science. 2019 Jun;364(6447):eaaw5181.

57. Baraduc P, R Duhamel J, Wirth S. Schema cells in the macaque hippocampus. Science. 2019 Feb;363(6427):635–9.

58. Knudsen EB, Wallis JD. Hippocampal neurons construct a map of an abstract value space. Cell. 2021 Sep;184(18):4640–50.e10.

59. Nieh EH, Schottdorf M, Freeman NW, Low RJ, Lewallen S, Koay SA, et al. Geometry of abstract learned knowledge in the hippocampus. Nature. 2021 Jul;595(7865):80–4.

60. Sosa M, Plitt MH, Giocomo LM. A flexible hippocampal population code for experience relative to reward. Nature Neuroscience. 2025 Jul;28(7):1497–509.

61. Vaswani A, Shazeer N, Parmar N, Uszkoreit J, Jones L, Gomez AN, et al. Attention is all you need. arXiv [csCL]. 2017 Jun.

62. Tampuu A, Matiisen T, Olafsdottir HF, Barry C, Vicente R. Efficient neural decoding of self-location with a deep recurrent network. PLoS Comput Biol. 2019 Feb;15(2):e1006822.

63. Zhang K, Ginzburg I, McNaughton BL, Sejnowski TJ. Interpreting neuronal population activity by reconstruction: unified framework with application to hippocampal place cells. J Neurophysiol. 1998 Feb;79(2):1017–44.

64. Schneider S, Lee JH, Mathis MW. Learnable latent embeddings for joint behavioural and neural analysis. Nature. 2023 May;617(7960):360–8.

65. Truccolo W, Eden UT, Fellows MR, Donoghue JP, Brown EN. A point process framework for relating neural spiking activity to spiking history, neural ensemble, and extrinsic covariate effects. J Neurophysiol. 2005 Feb;93(2):1074–89.

66. Pillow JW, Shlens J, Paninski L, Sher A, Litke AM, Chichilnisky EJ, et al. Spatiotemporal correlations and visual signalling in a complete neuronal population. Nature. 2008 Aug;454(7207):995–9.

67. Solla SA, Disterhoft JF, Wirtshafter HS. Conserved neural population geometry supports behavioral generalization. bioRxiv. 2026.

68. Jiang A, GoodSmith D, Ramirez-Matias J, Tortolani AF, Sheffield MEJ. Distinct CA3 inputs differentially shape the learning-dependent evolution of right CA1 spatial maps. Nat Commun. 2026 Apr;17(1):5682.

69. Carpenter J, Blackstad JS, Tingley D, Normand VA, Moser EI, Moser MB, et al. Investigating egocentric tuning in hippocampal CA1 neurons. Journal of Neuroscience. 2024 Sep;44(38):e0040242024.

70. Schmid P, Sesterhenn J. Dynamic Mode Decomposition of numerical and experimental data. In: 61st Annual Meeting of the APS Division of Fluid Dynamics. vol. 53. American Physical Society; 2008. .

71. Tu JH. Dynamic mode decomposition: Theory and applications. Princeton University; 2013.

72. Brunton BW, Johnson LA, Ojemann JG, Kutz JN. Extracting spatial–temporal coherent patterns in large-scale neural recordings using dynamic mode decomposition. Journal of Neuroscience Methods. 2016 Jan;258:1–15.

73. Prokop B, Gelens L. Data-driven discovery of dynamical models in biology. Nat Rev Phys. 2026 Jun;8(9):561–78.

74. Dotson NM, Yartsev MM. Nonlocal spatiotemporal representation in the hippocampus of freely flying bats. Science. 2021 Jul;373(6551):242–7.

75. Amaral DG, Witter MP. The three-dimensional organization of the hippocampal formation: a review of anatomical data. Neuroscience. 1989;31(3):571–91.

76. Stachenfeld KL, Botvinick MM, Gershman SJ. The hippocampus as a predictive map. Nature Neuroscience. 2017 Nov;20(11):1643–53.

77. George D, Rikhye RV, Gothoskar N, Guntupalli JS, Dedieu A, Ĺazaro-Gredilla M. Clone-structured graph representations enable flexible learning and vicarious evaluation of cognitive maps. Nat Commun. 2021 Apr;12(1):2392.

78. Raju RV, Guntupalli JS, Zhou G, Wendelken C, Lazaro-Gredilla M, George D. Space is a latent sequence: A theory of the hippocampus. Sci Adv. 2024 Aug;10(31):eadm8470.

79. Forli A, Yartsev MM. Understanding the neural basis of natural intelligence. Cell. 2024 Oct;187(21):5833–7.

80. Pastalkova E, Itskov V, Amarasingham A, Buzsaki G. Internally generated cell assembly sequences in the rat hippocampus. Science. 2008 Sep;321(5894):1322–7.

81. Kraus BJ, Robinson RJ 2nd, White JA, Eichenbaum H, Hasselmo ME. Hippocampal “time cells”: time versus path integration. Neuron. 2013 Jun;78(6):1090–101.

82. Omer DB, Las L, Ulanovsky N. Contextual and pure time coding for self and other in the hippocampus. Nature Neuroscience. 2023 Feb;26(2):285–94.

83. Sorscher B, Mel GC, Ocko SA, Giocomo LM, Ganguli S. A unified theory for the computational and mechanistic origins of grid cells. Neuron. 2023 Jan;111(1):121–37.e13.

84. Cueva CJ, Wei XX. Emergence of grid-like representations by training recurrent neural networks to perform spatial localization. In: International Conference on Learning Representations; 2018. .

85. Banino A, Barry C, Uria B, Blundell C, Lillicrap T, Mirowski P, et al. Vector-based navigation using grid-like representations in artificial agents. Nature. 2018 May;557(7705):429–33.

86. Whittington JCR, Warren J, Behrens TEJ. Relating transformers to models and neural representations of the hippocampal formation. In: International Conference on Learning Representations; 2022. .

87. Ramsauer H, Schäfl B, Lehner J, Seidl P, Widrich M, Adler T, et al. Hopfield Networks is All You Need. arXiv [csNE]. 2020 Jul.

88. Krotov D, Hopfield J. Large Associative Memory Problem in Neurobiology and Machine Learning. In: The Ninth International Conference on Learning Representations; 2021. .

89. Krenn M, Pollice R, Guo SY, Aldeghi M, Cervera-Lierta A, Friederich P, et al. On scientific understanding with artificial intelligence. Nat Rev Phys. 2022 Oct;4(12):761–9.

90. Lin Z, Akin H, Rao R, Hie B, Zhu Z, Lu W, et al. Evolutionary-scale prediction of atomic-level protein structure with a language model. Science. 2023 Mar;379(6637):1123–30.

91. Wang H, Fu T, Du Y, Gao W, Huang K, Liu Z, et al. Scientific discovery in the age of artificial intelligence. Nature. 2023 Aug;620(7972):47–60.

92. Brixi G, Durrant MG, Ku J, Naghipourfar M, Poli M, Sun G, et al. Genome modelling and design across all domains of life with Evo 2. Nature. 2026 Mar:1–13.

93. Doshi-Velez F, Kim B. Towards A Rigorous Science of Interpretable Machine Learning. arXiv [statML]. 2017 Feb.

94. Olah C, Cammarata N, Schubert L, Goh G, Petrov M, Carter S. Zoom In: An Introduction to Circuits. Distill. 2020.

95. Lindsey J, Gurnee W, Ameisen E, Chen B, Pearce A, Turner NL, et al. On the Biology of a Large Language Model. Transformer Circuits Thread. 2025. Available from: https://transformer-circuits.pub/2025/attribution-graphs/biology.html.

96. Ameisen E, Lindsey J, Pearce A, Gurnee W, Turner NL, Chen B, et al. Circuit Tracing: Revealing Computational Graphs in Language Models. Transformer Circuits Thread. 2025. Available from: https://transformer-circuits.pub/2025/attribution-graphs/methods.html.

97. de Vries SEJ, Lecoq JA, Buice MA, Groblewski PA, Ocker GK, Oliver M, et al. A large-scale standardized physiological survey reveals functional organization of the mouse visual cortex. Nature Neuroscience. 2020 Jan;23(1):138–51.

98. International Brain Laboratory, Angelaki D, Benson B, Benson J, Birman D, Bonacchi N, et al. A brain-wide map of neural activity during complex behaviour. Nature. 2025 Sep;645(8079):177–91.

99. MICrONS Consortium. Functional connectomics spanning multiple areas of mouse visual cortex. Nature. 2025 Apr;640(8058):435–47.

100. Hoffmann J, Borgeaud S, Mensch A, Buchatskaya E, Cai T, Rutherford E, et al. Training Compute-Optimal Large Language Models. arXiv [csCL]. 2022 Mar.

101. Jaegle A, Borgeaud S, Alayrac JB, Doersch C, Ionescu C, Ding D, et al. Perceiver IO: A General Architecture for Structured Inputs & Outputs. In: International Conference on Learning Representations; 2022. .

102. Reed S, Zolna K, Parisotto E, Colmenarejo SG, Novikov A, Barth-Maron G, et al. A Generalist Agent. arXiv [csAI]. 2022 May.

103. Brown T, Mann B, Ryder N, Subbiah M, Kaplan JD, Dhariwal P, et al. Language Models are Few-Shot Learners. Advances in Neural Information Processing Systems. 2020;33:1877–901.

104. Wei J, Tay Y, Bommasani R, Raffel C, Zoph B, Borgeaud S, et al. Emergent abilities of large language models. Transact Mach Learn Res. 2022 Jun.

105. Pachitariu M, Sridhar S, Pennington J, Stringer C. Spike sorting with Kilosort4. Nat Methods. 2024 May;21(5):914–21.

106. Zhang Y, Wang Y, Jimenez-Beneto DM, Wang Z, Azabou M, Richards BA, et al. Towards a ”Universal Translator” for Neural Dynamics at Single-Cell, Single-Spike Resolution. In: The Thirty-eighth Annual Conference on Neural Information Processing Systems; 2024. .

107. Loshchilov I, Hutter F. Decoupled weight decay regularization. arXiv [csLG]. 2017 Nov.

108. van Beest EH, Bimbard C, Fabre JMJ, Dodgson SW, Takacs F, Coen P, et al. Tracking neurons across days with high-density probes. Nat Methods. 2025 Apr;22(4):778–87.

